# A Dual-Pathway Prediction Error Model of Schizophrenia Spectrum Disorders: Bridging NMDA Hypofunction and Dopaminergic Hyperfunction

**DOI:** 10.64898/2026.01.28.702194

**Authors:** Shunsuke Sato, Tadahumi Kato, Taro Toyoizumi

## Abstract

Schizophrenia spectrum disorders (SSDs) present a profound clinical enigma, manifesting as a heterogeneous continuum ranging from the chaotic volatility of acute psychosis to the impenetrable rigidity of systematized delusions. While neurobiological research has independently implicated NMDA receptor hypofunction or dopaminergic hyperfunction as cardinal pathophysiological distinct mechanisms, a computational framework capable of bridging these distinct cellular deficits to the spectrum’s vast phenomenological diversity remains elusive. Here, we propose a biologically plausible neural model using a dynamic Bayesian inference with separable positive and negative prediction-error pathways. We demonstrate that NMDA hypofunction selectively blunts negative prediction errors, fostering rigid, bias-dominated beliefs, while dopaminergic hyperfunction uniformly amplifies error signals, driving volatile, observation-dominated states. Their interaction reconstructs SSDs as a continuous bias-volatility spectrum, accounting for key neurophysiological markers and offering a theoretical foundation for mechanism-based patient stratification.

## 2. Introduction

Generally, SSDs are defined by positive symptoms such as delusions and hallucinations, negative symptoms such as affective flattening and reduced motivation, and widespread cognitive impairment [1]. In addition to these clinical features, a substantial body of evidence has accumulated at the biological and network levels; for example, convergent biological data point to NMDA-receptor hypofunction on inhibitory interneurons and dysregulated dopaminergic signaling [2-4]. At the network level, empirical studies report attenuated mismatch negativity (MMN) or impaired predictive semantic processing, each tied to prediction-error signaling [5-7]. These common observations underlying SSDs are actively being studied to elucidate their characteristic pathophysiology.

Beyond their general characteristics, SSDs can exhibit a range of distinct symptoms across the spectrum [8]. The 11th revision of the International Classification of Diseases (ICD-11) defines the diagnostic categories included in SSDs, which encompass schizophrenia (SZ), acute transient psychotic disorder (ATPD), and delusional disorder (DD) [9]. While these categories are defined in ICD-11, the DSM criteria differ, and the diagnostic boundaries remain ambiguous. This lack of clarity persists because the biological basis for prioritizing specific symptoms remains unknown. Among these, SZ is one of the principal components. ATPD has an abrupt onset with rapidly fluctuating psychotic symptoms and usually remits over a short period. DD is characterized primarily by fixed, systematized delusions, and its remission rate with antipsychotic treatment is lower than that for SZ [10, 11]. Although clinical features of each disorder are increasingly well characterized, cross-diagnostic comparisons of their biological and network-level substrates—and the neurobiological mechanisms that give rise to the spectrum—remain scarce. These limitations highlight the difficulty of elucidating spectrum-level mechanisms solely through empirical observation.

Synthesizing individual findings into a coherent picture of the disorder benefits from theoretical frameworks that can propose algorithms underlying the overall pathophysiology. Current theories formalize SSD abnormalities as circuit breakdowns or errors in probabilistic inference [12–17]. These models have explained gamma-band activity abnormalities and working-memory impairment based on NMDA receptor hypofunction, cognitive deficits due to dopaminergic dysregulation, and hallucinations and delusions due to developmental abnormalities of attractor networks. Other accounts invoke aberrant salience attribution [16] or circular inference arising from an imbalance between excitation and inhibition [17]. More recently, predictive-coding frameworks have been used to reproduce MMN attenuation or hallucinatory-delusional symptoms by assuming abnormalities in precision control [17]. However, no existing model unifies the spectrum of SSD heterogeneity.

Because cross-diagnostic empirical studies are inherently constrained in linking human clinical observations to molecular mechanisms and to neural information processing, we adopt a theoretical approach to spectrum research. We present a neural circuit model of predictive-coding networks that implements the Kalman filter—an optimal algorithm for estimating latent states from sequential streams of noisy observations. In our model, this algorithm is realized by distinct positive and negative prediction-error (PE) pathways. Under this mapping, NMDA-receptor hypofunction in inhibitory interneurons reduces negative PE and increases positive PE, thereby biasing updates toward overestimating salient events and promoting delusional persistence. Moreover, upregulation of dopaminergic tone enhances PE gain, leading to the formation of unstable delusions by increasing sensitivity to sensory observations. This integrated mechanistic framework not only links cellular perturbations to neurophysiological markers and clinical symptoms within a unified model but also shows that the interaction of NMDA-bias and dopaminergic gain reproduces the heterogeneity of delusions across SZ, ATPD, and DD, yielding testable predictions.

## 3. Results

We instantiated a biologically plausible dual-pathway predictive-coding architecture implemented as a Kalman filter with separable positive and negative PEs streams. We modeled spectrum-wide symptom variation as parametric changes in two impairment parameters: NMDA-receptor function on inhibitory interneurons and dopaminergic tone. Empirical evidence supports two distinct neuronal populations, one encoding positive PE and the other encoding negative PE, in visual cortex [18], auditory cortex [19], midbrain [20], and prefrontal cortex [21], suggesting the dual-pathway organization as a general principle for neural coding. We assume that NMDA abnormalities bias the balance between the two pathways. In addition, we assume that abnormalities in dopaminergic tone disrupt the gain control of PEs. Converging animal and human studies indicate that dopamine modulates sensory precision and discrimination performance, consistent with its gain-control role [22]. Kalman filtering provides a normative account of sensory inference. The environment is described by hidden states that evolve over time and generate noisy observations. Because hidden states are not directly observable, the agent maintains a probabilistic estimate and updates it sequentially by combining the predicted prior of the model with current sensory evidence.

The model comprises a state (dynamics) equation and an observation (measurement) equation. To discuss biological correspondence simply, we present a one-dimensional Kalman filter model [23]. Let *h*_*t*_ denote the hidden state and *o*_*t*_ the observation at time *t*. For instance, *h* could represent the model of the frequency of a tone and *o* represent the firing rate of a sensory neuron. The state and observation equations are described by *h*_*t*_ = *a h* _*t*−1_ + *b* ξ_*t*_ and *o*_*t*_ = *c h*_*t*_ + *d* η_*t*_, respectively, where *a, b, c, d* are coefficients and ξ_*t*_ and η_*t*_ are standard Gaussian noises. The agent computes the Gaussian posterior distribution *p*(*h*_*t*_ |*o*_1:*t*_) of the hidden state *h*_*t*_ given the history of observations *o*_1:*t*_ = {*o*_1_, *o*_2_, …, *o*_*t*_}, parametrized by mean *M*_*t*_ and steady-state variance *V*. The variance is obtained by solving the discrete algebraic Riccati equation [23], written concisely as *V* = (*I* − *c K*)*V*_0_ with the Kalman gain *K* = *c V*_0_/(*c*^2^*V*_0_ + *d*^2^) and a priori variance *V*_0_ = *a*^2^*V* + *b*^2^. Then, the Kalman *K* = *c V*^0^ /(*c*^2^ *V*^0^+ *d*^2^) and a priori variance. *V*^0^= *a*^2^ *V* + *b*^2^ Then, the Kalman update equation of the mean is *M*_*t*_ = *a M*_*t*−*1*_+ *K*(*o*_*t*_ − *c a M*_*t*−*1*_). In this equation, the term *o*_*t*_ − *c a M*_*t*−*1*_ is the PE—the difference between the current observation and the predicted observation—and *K* is the Kalman gain, which determines how much the PE updates the state estimate. Our hypothesis is that excitatory-inhibitory circuits in the brain compute this Bayesian inference via separable positive and negative PEs pathways. The positive and negative PEs are defined as 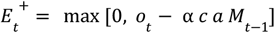 and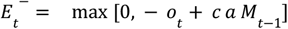, respectively. Specifically, the positive PE 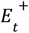 computes only the portion where the observation exceeds the prediction (the positive part of the observation *o*_*t*_ minus the prediction), and vice versa, for the negative PE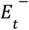. This separation of positive and negative PEs is achieved by adding the observation and prediction with opposite signs. The Kalman update rule for our dual-pathway model is described as

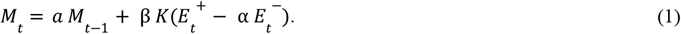

The coefficients α and β represent the NMDA-dependent inhibitory-bias parameter and dopamine-dependent gain-modulation parameter, respectively. While α = β = 1 is optimal for the Bayesian computation, we hypothesize this is not the case in SSDs. We investigate NMDA hypofunction (α < 1) and dopamine hyperfunction (β > 1) in the following sections. Note that in the equation for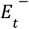, the sensory observation term − *o*_*t*_ is not multiplied by the inhibitory-bias factor α. This reflects empirical findings that NMDA receptor hypofunction is pronounced in frontal and temporal association cortices, while primary sensory areas remain relatively intact [24, 25]. Another issue is what sensory changes correspond to the positive or negative PE directions. As a rule of thumb, we assume that the rarer and more statistically salient error direction is encoded by the positive PE pathway to support sparse coding (see Discussion).

Equation (1) maps onto the cortical circuit model shown in Fig. 1. The biological interpretation is as follows. State-estimation units in deep cortical layers [26, 27] represent the estimated mean *M*.

**Fig. 1:**
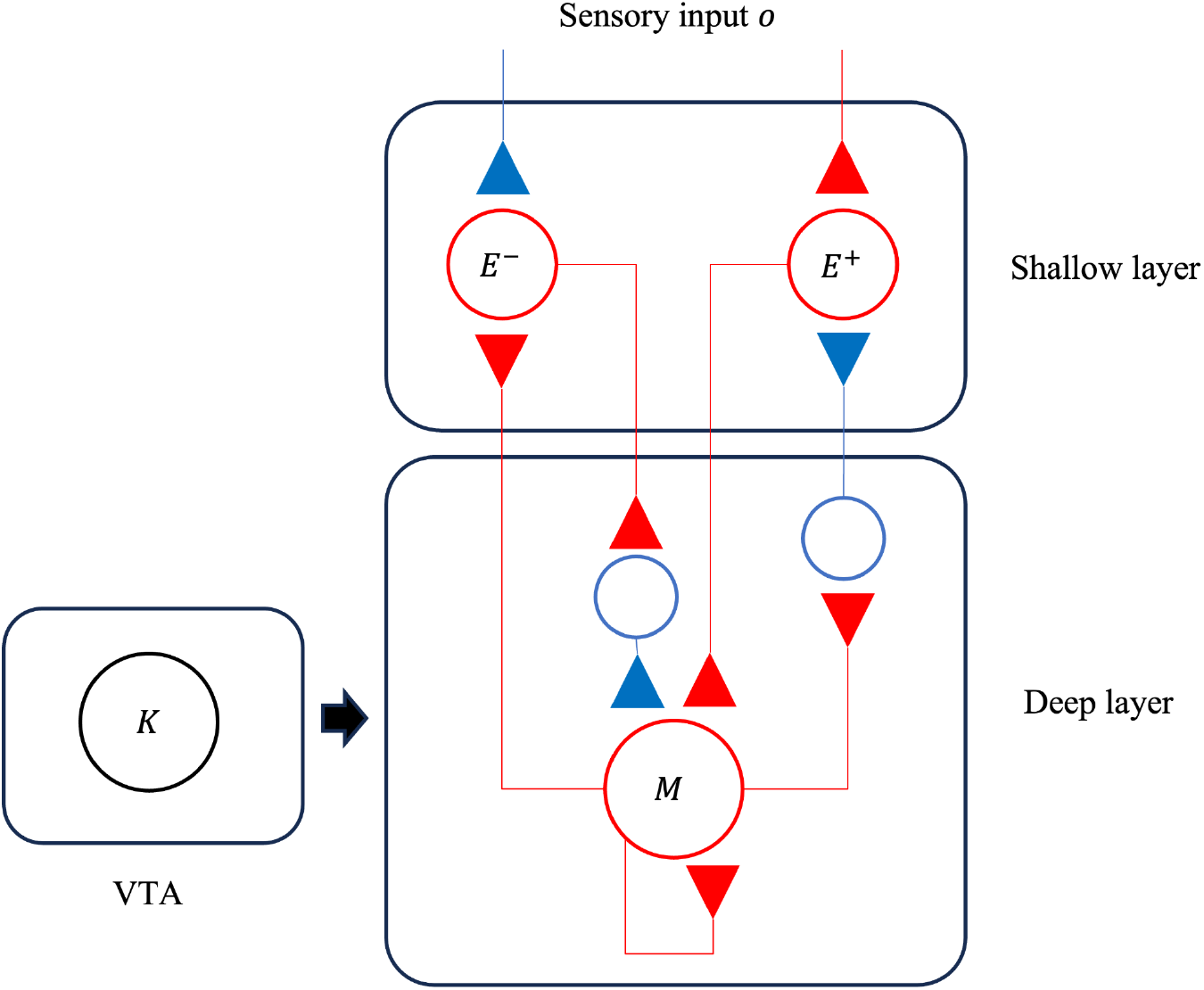
Cortical Circuit Implementation of the Dual-Pathway Prediction Error Model. The prediction error units *E*^±^ receive positive/negative top-down output from the previous time step’s state estimation unit *M*_*t*−*1*_,as well as negative/positive input from the observation *o*_*t*_. The prediction error units compute positive and negative prediction errors by summing the prediction input and the observation input with opposite signs. The factor *K* modulates the prediction error output. Using the adjusted prediction error output and the recurrent input *M*_*t*−*1*_, the state estimation unit *M*_*t*−*1*_ is updated to *M*_*t*_.

Positive and negative PEs units in shallow layers represent *E*^+^ [18]. A gain-control factor—putatively mediated by neuromodulatory mechanisms such as dopamine—encodes the Kalman gain *K*. The recurrent input weight to the state-estimation unit is denoted by *a*. The input weight from the state-estimation unit to the prediction-error units is *ac*. For simplicity of calculation, internal model parameters were fixed at *a* = 0. 9, *c* = 1, *d* = 1 and *b* = 0. 1. This parameter setting, where the system noise *b* is smaller than the observation noise *d*, is consistent with the setting of the previous Bayesian models of perception [28-30], which assume the external world evolves smoothly while sensory observations are noisy. The value of parameter $a$ was set to be less than 1 to ensure a stable system. Since small variations in these coefficients do not introduce an essential change to the simulation results, they were fixed throughout this study.

### Modeling NMDA Receptor Hypofunction: Bias dysfunction

Some research suggests that hallucinations and delusions in SSDs stem from an overweighting of prior beliefs relative to sensory evidence [8, 10], a phenomenon often termed “Strong Priors” [1, 5]. This mechanism was demonstrated in a Pavlovian conditioning task where participants learned to associate a visual cue with a 1000 Hz tone [31]. Patients with SSDs were significantly more likely to hallucinate the tone when presented with the visual cue alone (Fig. 2a). Furthermore, reduced MMN has been associated with more severe auditory hallucinations, and longitudinal evidence further suggests that baseline MMN deficits predict subsequent worsening of auditory hallucinations, implicating impaired PE processing in the maintenance or exacerbation of hallucinations [32, 33].

**Fig. 2:**
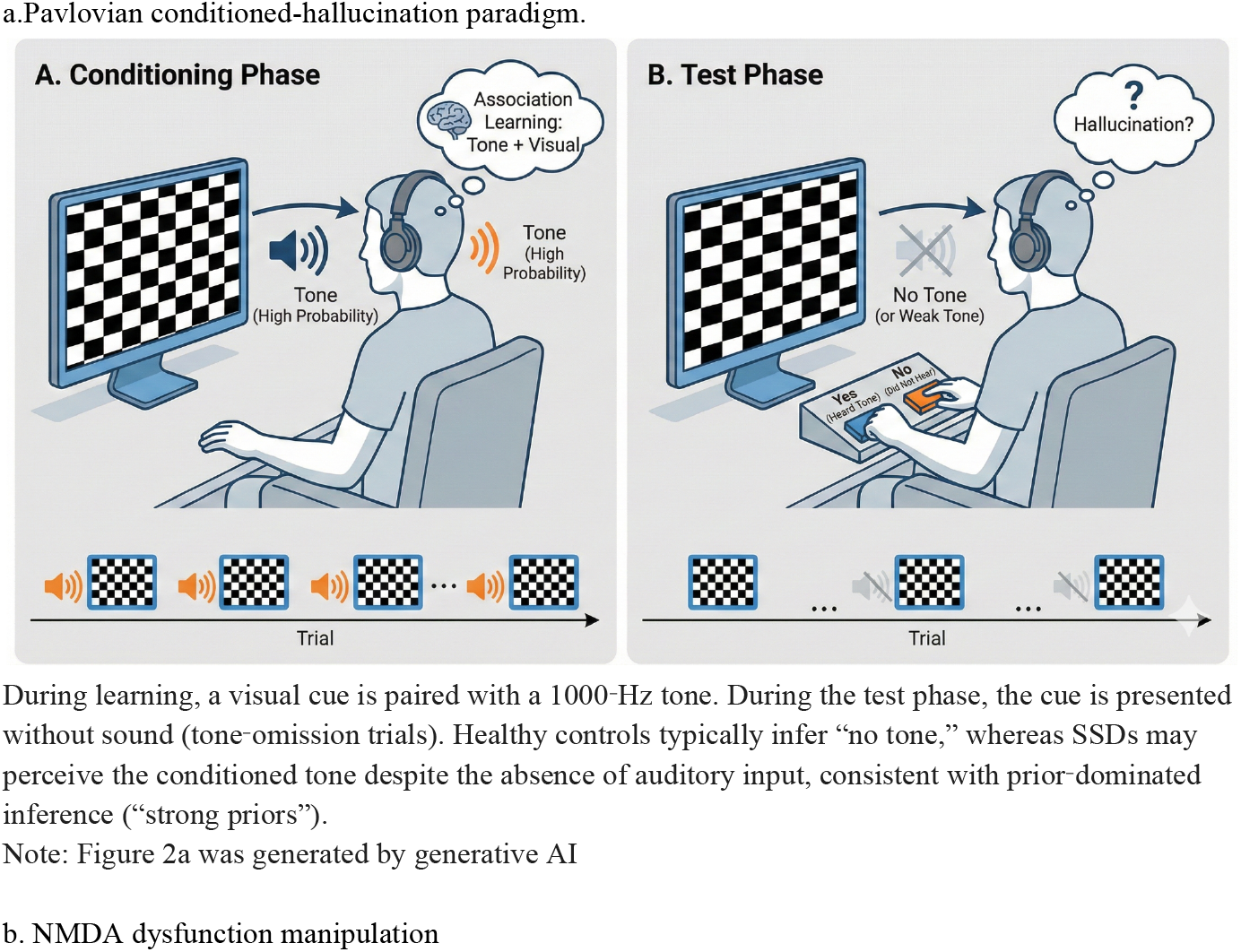
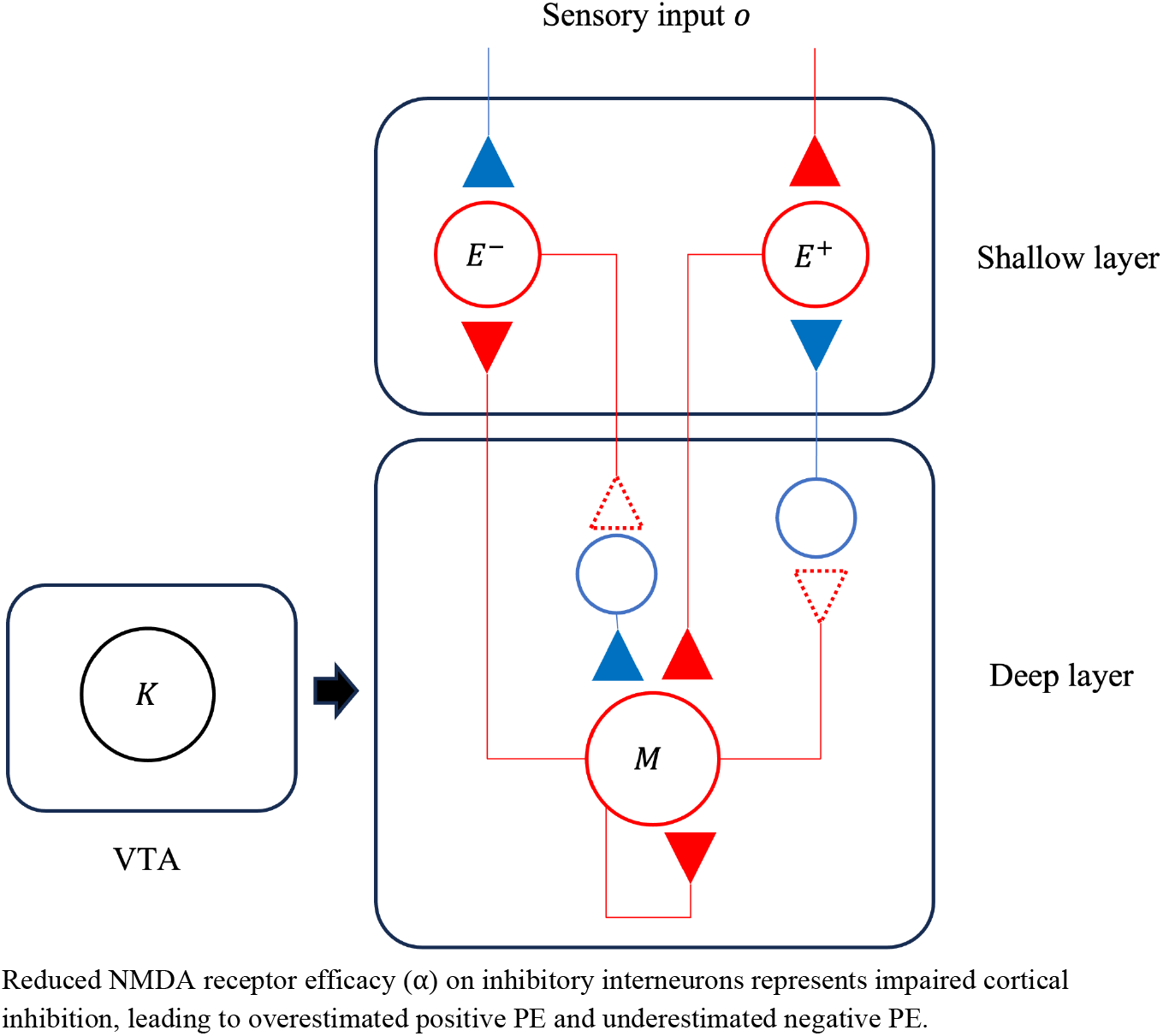
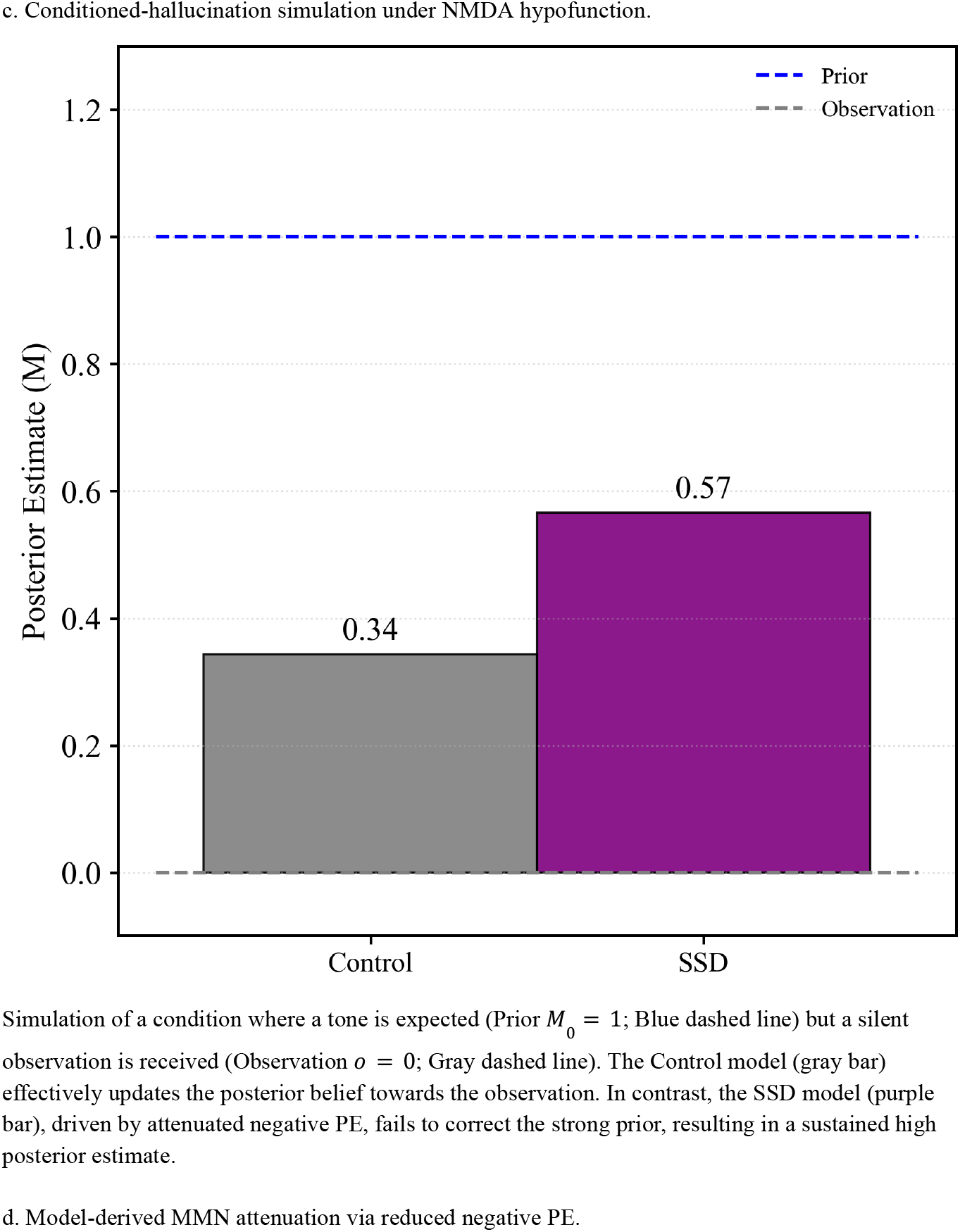
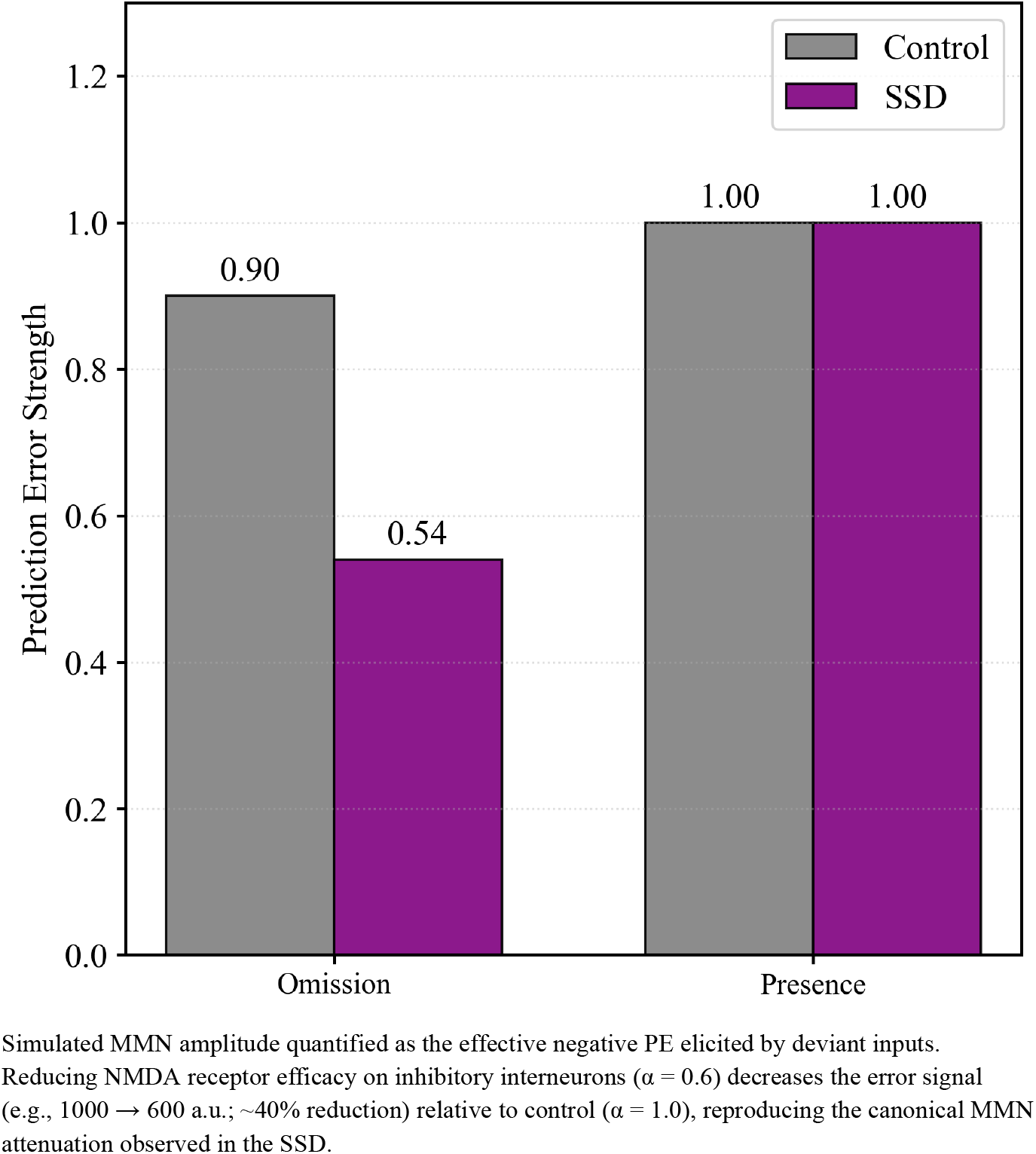
NMDA Receptor Hypofunction-Induced Bias Dysfunction: A Unified Mechanism for Hallucinations and MMN Attenuation.

We replicated these findings using our dual-pathway PE model, interpreting both conditioned hallucinations and MMN reduction as consequences of imbalance in the PE pathways. First, to simulate conditioned hallucinations, we defined the hidden state *h* as the volume of a 1000 Hz tone and the observation *o* as the auditory input (sensory firing rate). We assumed that the auditory input increases with the stimulus volume. We set a strong prior expectation of a tone (*M*_0_ = 1) but provided no sensory input (*o* = 0). In the control condition (α = 1), the discrepancy between prediction and observation generates a large negative PE, which drives the posterior estimate significantly towards the observation, correctly inferring the absence of the high-frequency tone (Fig.2c). However, in the SSD condition, we set α = 0. 6, reflecting the biological finding of a 30–50% reduction in NMDA receptor function at inhibitory dendrites [2-4] (Fig. 2b). Consequently, the belief update is blunted, and the posterior estimate remains abnormally close to the prior (Fig. 2c). This “rigidity of belief” represents a hallucination: the internal expectation of a high pitch persists, consistent with the Bias Against Disconfirmatory Evidence [1], which posits that patients with schizophrenia tend to ignore information that contradicts their existing beliefs.

Second, using the same setup, we simulated MMN (Fig. 2d). In this context, *h* and *o* similarly represent the volume and sensory input of a standard tone or of a deviant tone. MMN is electrophysiologically defined as the sum of neural responses to the omission of the standard tone, which produces a negative PE, and the presence of the deviant tone, which produces a positive PE. The NMDA hypofunction directly reduces the negative PE by a factor α, but does not appreciably increase the positive PE because the deviant tone was previously unexpected (i.e., α *M*_0_ = 0 regardless of the value of α). Thus, the model shows that the same mechanism—attenuated negative PE—simultaneously explains the “persistence of hallucinatory beliefs” and the “reduction of MMN”, offering a unified mechanistic account for these correlated clinical features. Note that while we modeled the above specific population, the actual brain contains diverse populations, including those where the response decreases linearly with increasing volume. For such populations, an Omission acts as a “Presence” (generating intact positive PE), while a Presence acts as an “Omission” (generating attenuated negative PE). Consequently, the aggregate MMN, summing across these diverse populations, is predicted to be attenuated regardless of whether the deviation involved an omission or a presence.

### Modeling Dopaminergic Hyperfunction: Gain dysfunction

In contrast to the “overweighting of priors” described above, SSDs paradoxically exhibit an “overweighting of sensory evidence” in perceptual domains. Patients often show resistance to visual illusions, perceiving the true size or shape of objects more accurately than healthy controls because contextual priors exert less influence. A prominent example is the Ebbinghaus illusion, where the apparent size of a central disk is biased by surrounding flankers; individuals with SSDs display reduced susceptibility to this illusion [34, 35], an effect that can be restored by dopamine-blocker treatment. In this simulation, we reproduced this “overweighting of observation” as a consequence of dopaminergic hyperfunction. Converging PET meta-analytic and pharmacological-challenge data indicate that striatal dopaminergic release is increased by about 15% in SSDs [36]. We operationalized this as a 15% increase of the gain modulator β that scales up the Kalman-gain (Fig. 3a) with the inhibitory coefficient set to its baseline value of 1.0. To evaluate perceptual consequences, we simulated the Ebbinghaus illusion. While there are multiple explanations for this size-contrast effect, a representative mechanism is lateral inhibition: when the surrounding circles are large (or small), lateral inhibition makes the central circle appear smaller (or larger) [37]. We modeled this illusion-inducing context as a biased prior expectation under two opposing conditions:

**Fig. 3:**
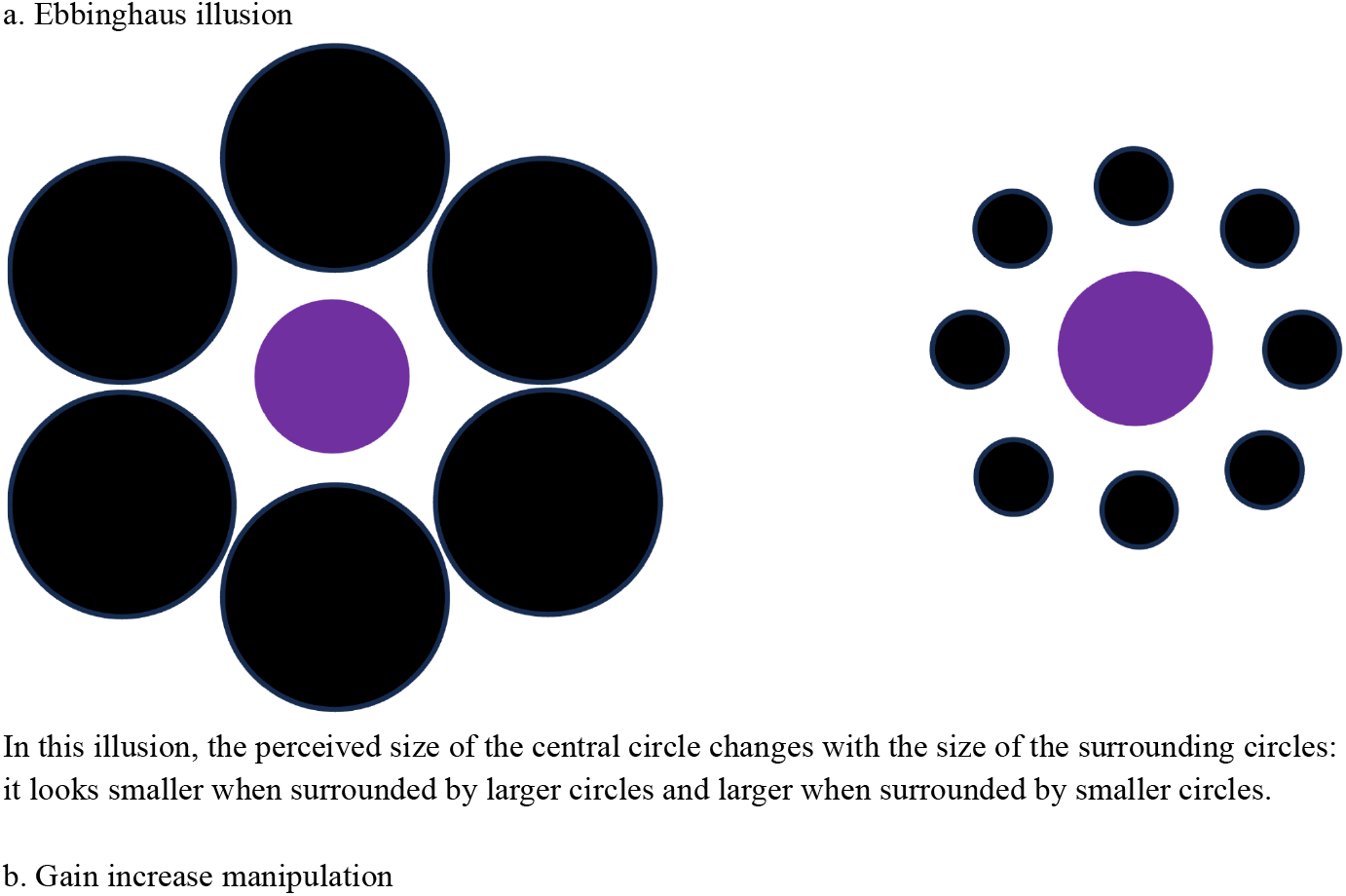
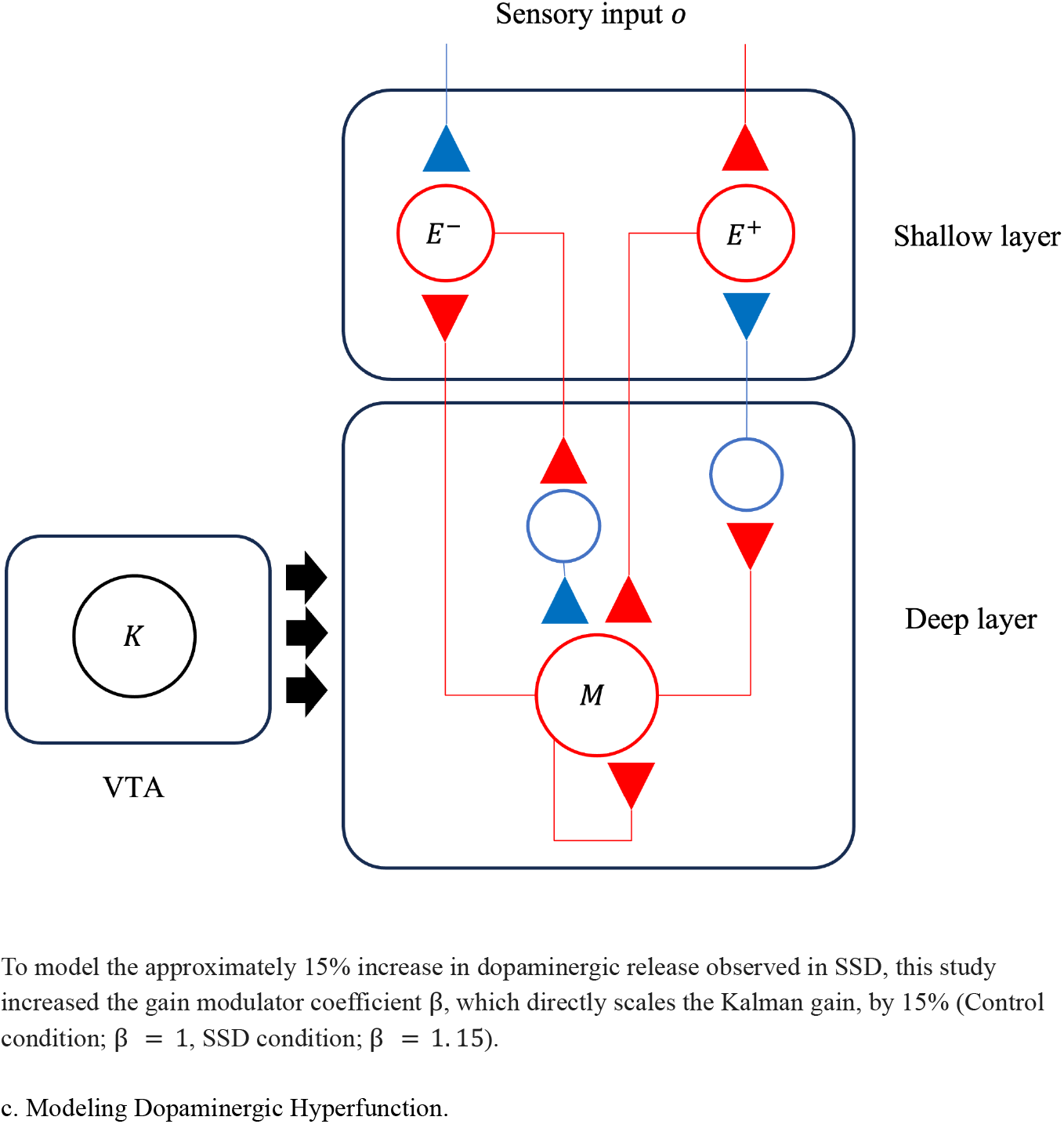
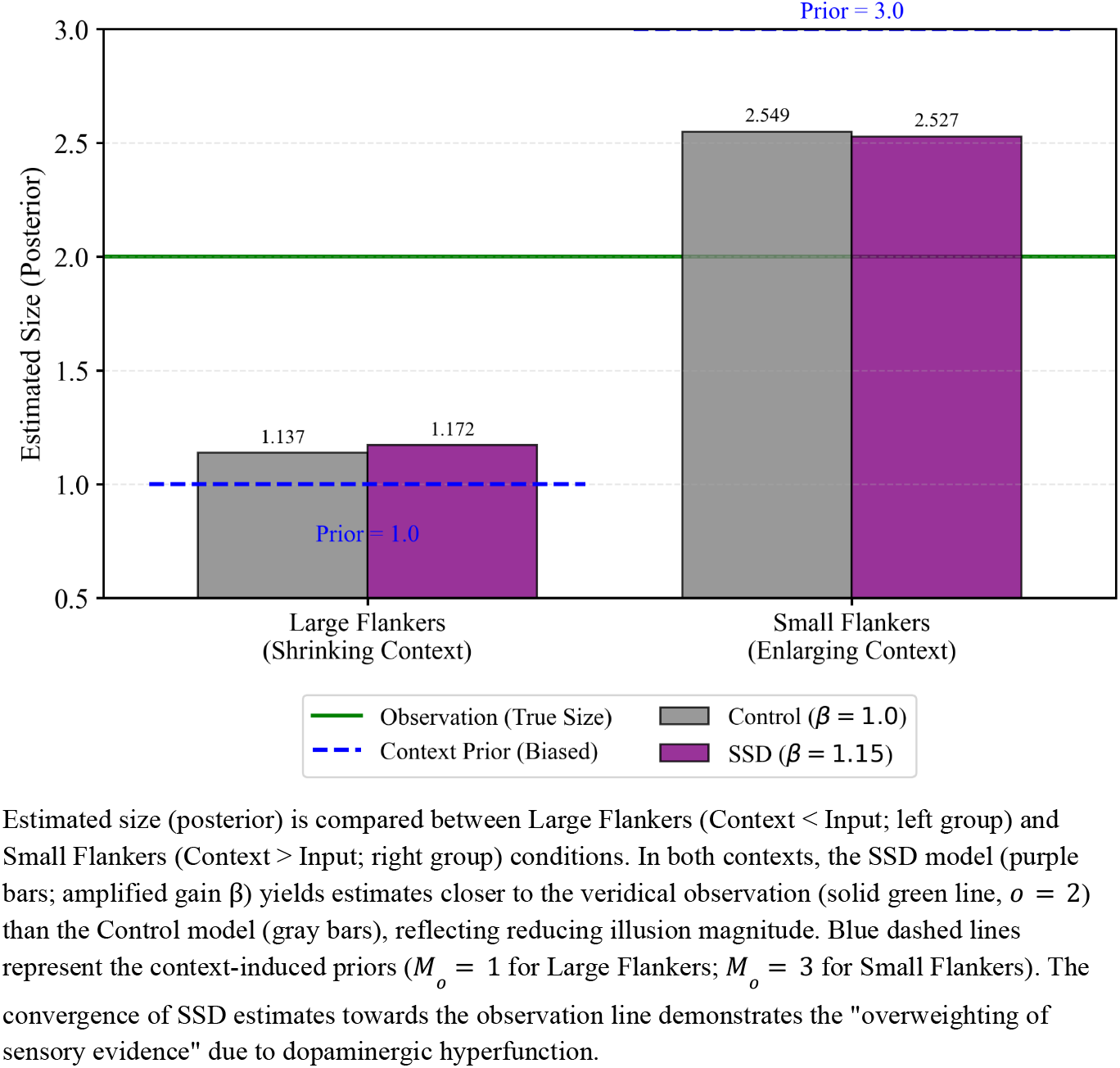
Dopaminergic Hyperfunction-Induced Gain Dysfunction: Overweighting of Sensory Evidence and Reduced Illusion Susceptibility.

1. Large Flankers (Context < Input): Large surrounding circles create a size-contrast effect, causing the central disk to appear smaller than it physically is. We modeled this illusion-inducing context as a biased prior expectation set to *M*_0_ = 1 (smaller expectation), relative to the true sensory input of *o* = 2.
2. Small Flankers (Context > Input): Small surrounding circles create a contrast effect making the central disk appear larger. We modeled this by setting the context prior to *M*_0_= 3 (larger expectation), relative to the true sensory input of *o* = 2.

In this illusion setting, inference was single-shot (*t*=1): the central disk drives *o* = 2 (normalized units), and the flankers provided a prior estimate biased by the context (*M*_0_ = 1 for Large Flankers, *M*_0_ = 3 for Small Flankers). Here, *h* represents the visual object size, and *o* represents the firing rate of visual neurons selectively responsive to size. We assume a population of neurons where the firing rate increases linearly with object size. Note that the visual cortex likely contains neurons tuned to both “larger” and “smaller” sizes; however, the core computational mechanism—where dopaminergic gain amplifies the sensory prediction error relative to the prior—holds regardless of whether the specific cell population is tuned to large or small objects. Therefore, we focus on modeling the cell population whose response increases linearly with the size of the central disk.

Mechanistically, this process is mediated by dopamine-enhanced PEs. The discrepancy between the true sensory size and the biased expectation generates a positive or negative PE. This error signal drives the update of the size estimate towards the true value. If the gain is standard, the update is partial, and the posterior estimate remains pulled towards the prior (illusion effect). However, under the SSD condition with increased dopamine, the weight of the sensory error is amplified. This demonstrates how dopaminergic gain control directly regulates the balance between sensory evidence and contextual priors. Note also that the positive PEs implies dependence on the inhibitory coefficient as well. This suggests that NMDA hypofunction could concurrently increase *E*^+^ effectively mimicking the effect of dopaminergic gains by boosting the error signal and reducing illusion susceptibility. This interaction introduces a directional asymmetry. Unlike the hallucination simulation where the absence of input (*o* = 0) made the attenuation of negative PE catastrophic (*M*_0_ = 1), in the illusion task (*o* = 2), both positive and negative PEs have substantial discrepancies. Yet, because NMDA hypofunction selectively attenuates the negative PE, the model predicts a subtle divergence: the correction of belief is less effective when the context overestimates the target (negative error) compared to when it underestimates it (positive error). This implies that the reduction in illusion susceptibility in SSDs may be arguably more pronounced for ‘shrinking’ contexts (which generate positive errors) than for ‘enlarging’ contexts.

The above simulations, the first of which modeled NMDA receptor hypofunction and the second, in addition to the NMDA hypofunction, dopaminergic dysregulation, reproduce neurophysiological characteristics of SSD from a molecular impairment basis. We next turn to delusional symptoms, the core clinical features. We asked whether our model can generate a spectrum of delusional dynamics from a mixture of NMDA-receptor hypofunction and dopaminergic dysregulation.

### Spectrum Reproduction of Delusional Dynamics via Parameter Variation

Clinical evidence demonstrates that patients with SSDs frequently overestimate others’ hostility [38, 39], a phenomenon closely associated with the emergence and maintenance of persecutory delusions. We modeled the intensity of another person’s hostility as a hidden state and the magnitude of their hostile behavior (e.g., facial expression or tone of voice) as the observation. In this framework, higher values of *h* correspond to greater underlying hostility, and higher values of *o* represent stronger

perceptible signals of threat. The simulation task was to estimate the changing level of hostility (*h*_*t*_) based on a sequence of noisy behavioral observations (*o*_*t*_). The true dynamics of hostility followed a standard linear-Gaussian process over (*t* = 30) steps (Fig. 4a). The observer adopted the same linear-Gaussian model, as described earlier, and computed posteriors via Kalman updating to track the latent hostility. We compared four parameter sets, Control (α = 1, β = 1), Dopamin hyperfunction ( α = 1, β = 1. 15), NMDA receptor hypofunction (α = 0. 6, β = 1) and Both (α = 0. 6, β = 1. 15). Under the Both condition, hostility was persistently overestimated relative to the Control condition, resembling fixed paranoid ideas seen in SZ. The Dopamine hyperfunction condition produced rapidly fluctuating estimates, reproducing the unstable delusions seen in ATPD. The NMDA receptor hypofunction condition overestimated hostility and maintained this elevation longer than the Both condition, modeling the rigid, poorly corrigible beliefs characteristic of DD (Fig. 4b). Temporal instability was quantified as the root mean square of successive differences (RMSSD) of the estimates (Fig. 4c, Method 5.1). The Dopamine hyperfunction condition showed higher RMSSD (more unstable delusions) than the Both condition, while the NMDA receptor hypofunction condition showed lower RMSSD (more stable delusions). A heat map (Fig. 4d) illustrates the parameter dependence of RMSSD.

**Fig. 4:**
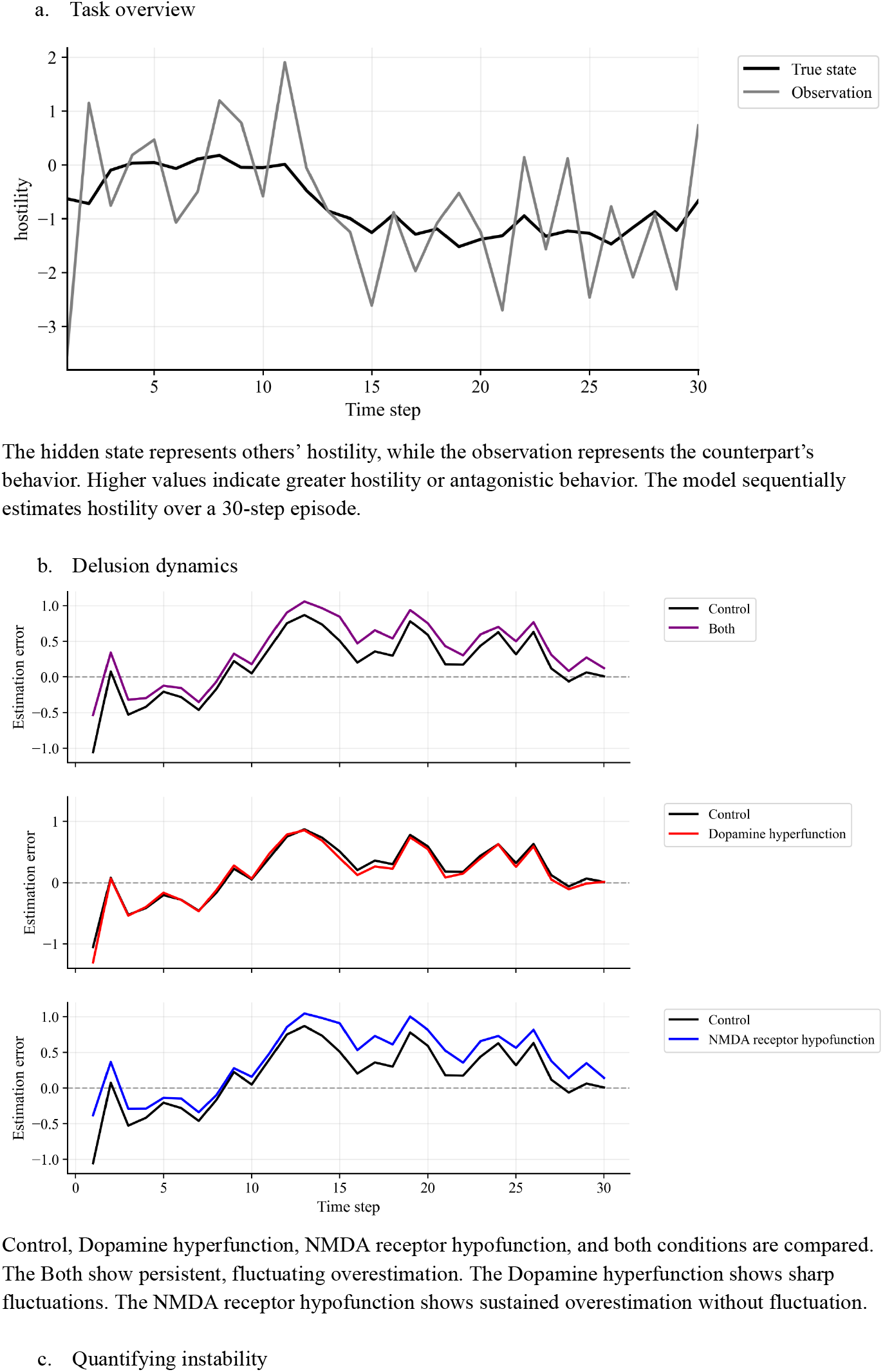
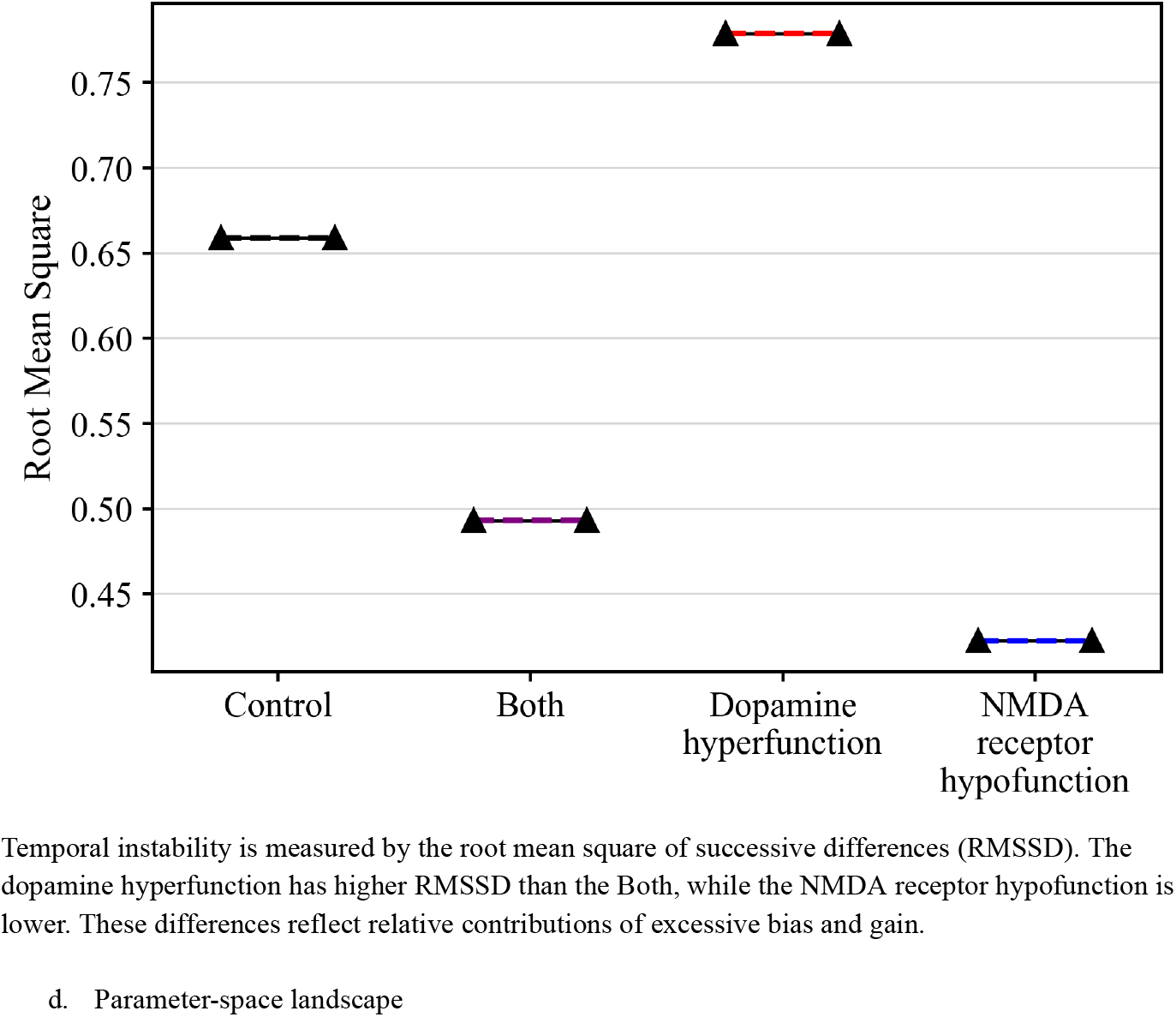
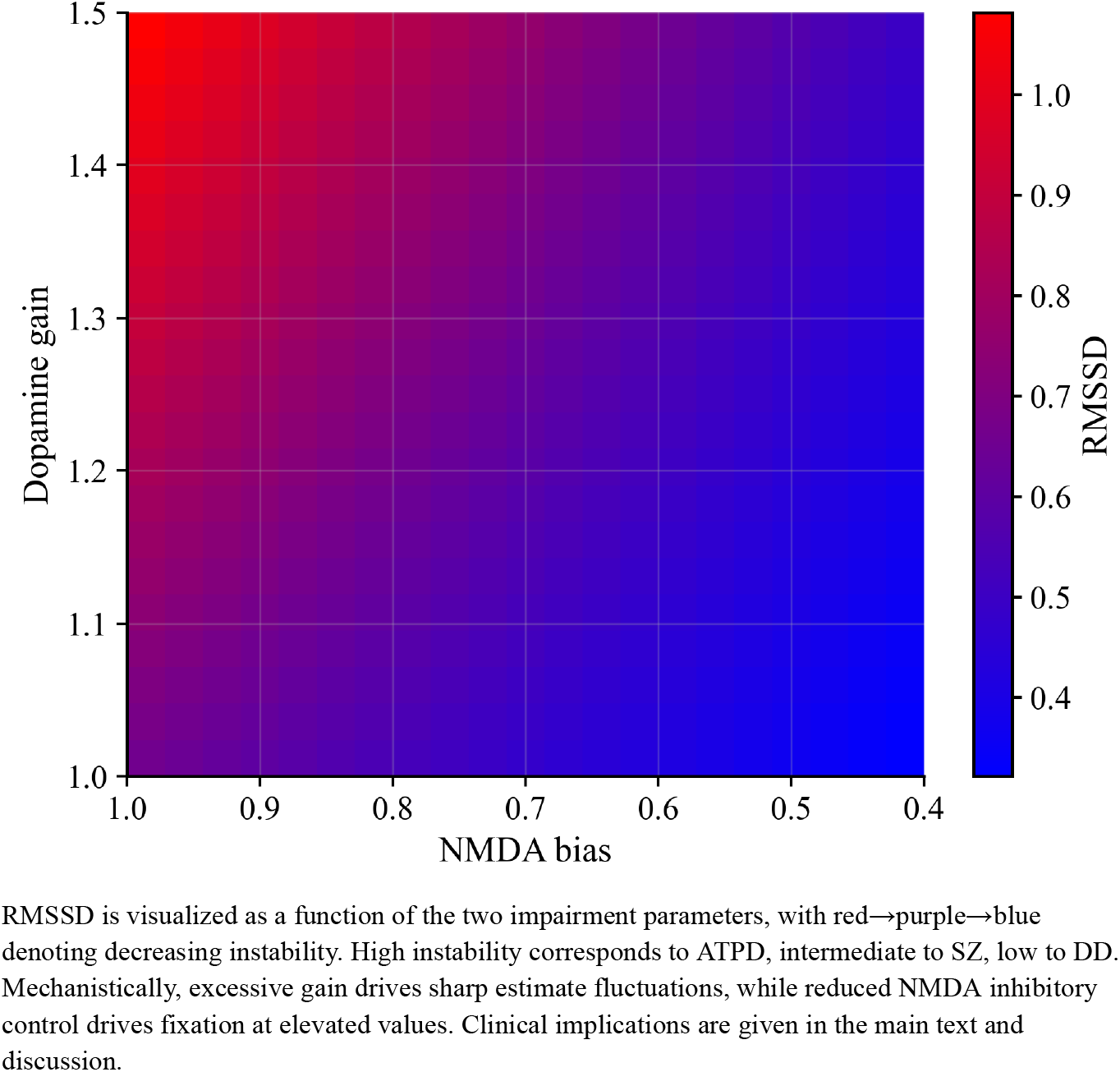
Spectrum Reproduction of Delusional Dynamics via Interaction of NMDA Bias and Dopaminergic Gain.

Reducing α decreases negative PE and increases positive ones, producing a pathological bias toward overestimation. Increasing β amplifies update amplitudes in both positive and negative PEs, generating volatile misestimation. Their interaction forms a two-dimensional bias–volatility plane, providing a quantitative spectrum linking disorder-specific delusional phenotypes.

## 4. Discussion

We present a two-path predictive-coding model that implements a linear Kalman filter with separable positive and negative PEs streams [18, 19-21, 23, 26] In this model, NMDA-receptor hypofunction on inhibitory interneurons maps to asymmetric error efficacy [2–4], and dopaminergic neuromodulation maps to gain control [22, 40, 41]. Through this mapping, the model links cellular alterations to neurophysiological markers and to spectrum-wide heterogeneity in delusional dynamics across SZ, ATPD, and DD [1, 5, 8-12, 42-44].

The present model assumes linear–Gaussian dynamics and a one-dimensional latent state and observation. However, delusional content and symptom trajectories are multidimensional and context-dependent. Delusions in SSDs are, in reality, not limited to persecutory delusions resulting in hostility as postulated in this study, but include other types of delusions, such as grandiosity or jealousy delusions. Extending the model to multivariate and hierarchical latent dynamics would allow us to capture richer symptom mixtures and cross-domain fluctuations [26, 27]. Thus, generalization to nonlinear, hierarchical generative models will be necessary [26, 29]. We approximate dopamine as a pure gain controller and map NMDA hypofunction exclusively to inhibitory interneurons; other neuromodulators and cell classes, including glial cells, likely contribute to gain control and error computation [22, 46-50]. We focused on a single readout; broader, multimodal batteries across sensory, cognitive, and social inference domains would better constrain parameter estimates and would be an important direction for future work.

In terms of computational implementation, prior studies have derived neural implementations or approximations of Kalman filtering and related Bayesian computations in recurrent circuits and probabilistic population codes [28, 51, 52]. These approaches often include multidimensional formulations and relax the Gaussian assumption; however, they have not been widely applied to explain SSD phenomenology. In parallel, influential accounts of psychosis have sought to unify the apparent paradox of “overweighted priors” and “overweighted sensory evidence”. For instance, the circular inference model proposes that excitatory-inhibitory imbalance causes descending priors and ascending sensory inputs to reverberate, effectively being “double-counted” as both prior and evidence [53]. Similarly, hierarchical predictive coding accounts suggest that aberrant precision control at different hierarchy levels can lead to simultaneous sensory hypersensitivity (low-level) and rigid beliefs (high-level) [54, 55]. These models invoke recurrent processes that converge to specific fixed points (attractors) [13, 15–17, 26, 31], framing delusion formation largely as a static inference problem where the system settles into a stable, albeit aberrant, belief state. They do not, however, explicitly capture the temporal dynamics of how beliefs evolve and fluctuate in response to a continuously changing environment. In contrast, our linear–Gaussian implementation focuses on the sequential tracking of evolving latent states. This allows us to model the trajectory of belief updating, thereby linking physiological parameters not only to the static presence of delusions, but also to their temporal instability and persistence across the SSDs. Our model integrates these lines of work by applying an exact Kalman implementation to SSD phenomenology.

Within this framework, a critical question regarding the content of delusions is why negative, threat-related delusions are more prevalent than positive, benevolent ones, although our simulations instantiate a circuit that estimates others’ hostility [56]. This asymmetry can be understood through the principles of sparse coding and the statistical properties of natural stimuli. To maximize metabolic efficiency, neural systems employ sparse coding, where rare, high-information events drive active firing (excitation) while common, redundant background events are represented by silence or low activity [57]. Natural signals typically follow power-law distributions—such as the 1/*f* scaling of sound frequency or the scale-invariant distribution of visual object sizes—where high-magnitude events are statistically rare outliers [60-62]. If we posit that social signals follow similar statistics, where overt hostility is a “rare” deviation from a neutral baseline [59, 61, 62], then efficient coding dictates that hostility should be mapped to the excitatory (positive prediction error) channel.

Consequently, the specific impairment of this excitatory pathway—disinhibition due to NMDA receptor hypofunction—would selectively amplify these rare, threat-related signals, leading to the preferential formation of persecutory delusions [2–4, 56].

This mechanistic insight leads to the model’s most important empirically testable prediction: clinical stratification that directly enables treatment selection. Specifically, sign-selective prediction-error signaling can be probed using a sign-balanced oddball paradigm [63, 64]. In this task, deviations occur in opposite directions relative to the learned regularity, yielding neural error signatures for conditions dominated by positive and negative prediction errors [63, 64]. These responses decompose into two components mapping onto the model’s impairment dimensions. First, the overall error signal strength across both directions indexes global gain, which reflects dopaminergic amplification [22, 40]. Second, the directional asymmetry in responses indexes sign-specific error efficacy, reflecting NMDA-related bias via attenuated negative prediction errors [5, 65]. This framework predicts distinct patient profiles. Patients with strong overall error signals but little asymmetry likely fall into a gain-dominated regime. Conversely, patients with pronounced asymmetry likely fall into a bias-dominated regime. Such stratification guides actionable treatment selection. Bias-dominated individuals should respond preferentially to NMDA-receptor augmentation using agents like D-serine, glycine, or sarcosine [65-67]. In contrast, gain-dominated individuals should benefit more from dopaminergic modulation using typical or atypical antipsychotics [41].

Finally, the relevance of this framework may extend beyond the SSDs. Sign-selective prediction-error representations are observed across cortical and subcortical systems [18-21]. In SZ, where frontal dysfunction is prominent, disruption of sign-selective PE computations within prefrontal networks would be expected to bias inference toward threat [68]. The prefrontal cortex plays a central role in estimating the potential threat of external stimuli; thus, its dysfunction may lead to exaggerated threat inference, consistent with persecutory delusions [56, 69]. By the same logic, analogous disruptions in other nodes should yield symptom profiles that reflect the computations subserved by those regions. For example, if limbic circuits such as the amygdala and anterior cingulate implement sign-selective PE for affective signals, their dysregulation could contribute to affective lability as observed in bipolar disorder [70, 71]. These extensions remain speculative and go beyond the regimes directly simulated here.

## 5. Methods

### 5.1 Data Analysis

Belief trajectories in the hostility estimation task were analyzed using Bias (mean signed error) and Volatility (RMSSD). The Root Mean Square of Successive Differences (RMSSD) is defined as:

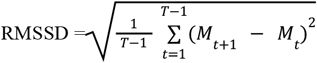

where *T* is the total number of time steps.

## 6. Data Availability

No datasets were generated or analyzed during the current study as it is a mathematical model.

## 7. Code availability

Custom code used for simulation is available in the attached Supplemental Material.

## 8. Author Contributions

S.S. and T.T. conceived the study, developed the computational model, performed the simulations, and wrote the manuscript. T.K. supervised the biological theoretical framework and facilitated the discussion, and contributed to the writing of the manuscript.

## Reference

1. Tandon, R., Keshavan, M.S. & Nasrallah, H.A. Schizophrenia, “just the facts”. Schizophrenia Research 110, 1–23 (2009).

2. Nakazawa, K., Zsiros, V., Jiang, Z., Nakao, K., Kolata, S., Zhang, S. & Belforte, J.E. GABAergic interneuron origin of SZ pathophysiology. Neuropharmacology 62, 1574–1583 (2012).

3. Billingslea, E.N., Tatard-Leitman, V.M., Anguiano, J., Jutzeler, C.R., Suh, J., Saunders, J.A., Morita, S. & Featherstone,, R.E. et al. Parvalbumin Cell Ablation of NMDA-R1 Causes Increased Resting Network Excitability with Associated Social and Self-Care Deficits. Neuropsychopharmacology 39, 1603–1613 (2014).

4. Kotzadimitriou, D., Nissen, W., Paizs, M., Newton, K., Harrison, P.J., Paulsen, O. & Lamsa, K. Neuregulin 1 Type I Overexpression Is Associated with Reduced NMDA Receptor-Mediated Synaptic Signaling in Hippocampal Interneurons Expressing PV or CCK. eneuro 5, ENEURO.0418-17.2018 (2018).

5. Garrido, M.I., Kilner, J.M., Stephan, K.E. & Friston, K.J. The mismatch negativity: A review of underlying mechanisms. Clinical Neurophysiology 120, 453–463 (2009).

6. Shelley, A., Ward, P., Catts, S., Michie, P., Andrews, S. & McConaghy, N. Mismatch Negativity: An Index of a Preattentive Processing Deficit in SZ. Biological Psychiatry 30, 1059–1062 (1991).

7. Sharpe, V., Weber, K. & Kuperberg, G.R. Impairments in Probabilistic Prediction and Bayesian Learning Can Explain Reduced Neural Semantic Priming in SZ. Schizophrenia Bulletin 46, 1558–1566 (2020).

8. Os, J.v., Linscott, R.J., Myin-Germeys, I., Delespaul, P. & Krabbendam, L. A systematic review and meta-analysis of the psychosis continuum: evidence for a psychosis proneness-persistence-impairment model of psychotic disorder. Psychological Medicine 39, 179–195 (2008).

9. Guloksuz, S. & Os, J.v. Waiting for the Funeral of “SZ” and the Baby Shower of the Psychosis Spectrum. Frontiers in Psychiatry 12, (2021).

10. González-Rodríguez, A. & Seeman, M.V. Differences between delusional disorder and SZ: A mini narrative review. World Journal of Psychiatry 12, 683–692 (2022).

11. Pillmann, F., Wustmann, T. & Marneros, A. Acute and transient psychotic disorders versus persistent delusional disorders: a comparative longitudinal study. Psychiatry and Clinical Neurosciences 66, 44–52 (2012).

12. Vierling-Claassen, D., Siekmeier, P., Stufflebeam, S. & Kopell, N. Modeling GABA alterations in SZ: a link between impaired inhibition and altered gamma and beta range auditory entrainment. Schizophrenia Research 102, 7 (2008).

13. Compte, A. Synaptic mechanisms and network dynamics underlying spatial working memory in a cortical network model. Cerebral Cortex 10, 910–923 (2000).

14. Cohen, J.D. & Servan-Schreiber, D. Context, cortex, and dopamine: a connectionist approach to behavior and biology in SZ. Cognitive Modeling, 377–422 (2002).

15. Hoffman, R.E. & Dobscha, S.K. Cortical pruning and the development of SZ: a computer model. Schizophrenia Bulletin 15, 477–490 (1989).

16. Kapur, S. Psychosis as a state of aberrant salience: a framework linking biology, phenomenology, and pharmacology in SZ. American Journal of Psychiatry 160, 13–23 (2003).

17. Jardri, R. & Denève, S. Circular inferences in SZ. Brain 136, 3227–3241 (2013).

18. Jordan, R. & Keller, G.B. Opposing Influence of Top-down and Bottom-up Input on Excitatory Layer 2/3 Neurons in Mouse Primary Visual Cortex. Neuron, (2020).

19. Parras, G.G., Nieto-Diego, J.L., Pérez-González, M.J., Halliday, N., López-Poveda, L.M., Covey, E. & Malmierca, M.S. Neurons along the auditory pathway encode a hierarchy of prediction errors. Nature Communications 8, (2017).

20. Matsumoto, M. & Hikosaka, O. Two types of dopamine neuron distinctly convey positive and negative motivational signals. Nature 459, 837–841 (2009).

21. Monosov, I.E. & Hikosaka, O. Regionally distinct processing of rewards and punishments by the primate ventromedial prefrontal cortex. The Journal of Neuroscience 32, 10318–10330 (2012).

22. Schwartenbeck, P., FitzGerald, T., Bossaerts, P., Huys, Q., Dolan, R.J. & Friston, K.J. The dopaminergic midbrain encodes the expected certainty about desired outcomes. Cerebral Cortex 25, 3434–3445 (2014).

23. Kalman, R.E. A New Approach to Linear Filtering and Prediction Problems. Journal of Basic Engineering 82, 35–45 (1960).

24. Akbarian, S., Sucher, N.J., Bradley, D., Tafazzoli, A., Trulson, D., Jones, E.G., Huntsman, M.M., Al-Ramahi, I., Parker, C.C. & Noble, E.P. Selective alterations in gene expression for NMDA receptor subunits in prefrontal cortex of schizophrenics. The Journal of Neuroscience 16, 19–30 (1996).

25. Mathew, S.V., Law, A.J., Lipska, B.K., Weinberger, D.R., Harrison, P.J. & Reynolds, G.P. A Novel Abnormality in the N-Methyl-D-Aspartate Glutamate Receptor: A Study of GluN2B Protein and Binding Density in the Human Hippocampus in Schizophrenia. Neuroscience Letters 304, 9–12 (2001).

26. Bastos, A.M., Usrey, W.M., Adams, R.A., Mangun, G.R., Fries, P. & Friston, K.J. Canonical microcircuits for predictive coding. Neuron 76, 695–711 (2012).

27. Felleman, D.J. & Essen, D.C.V. Distributed hierarchical processing in the primate cerebral cortex. Cerebral Cortex 1, 1–47 (1991).

28. Wilson, R.C. & Finkel, L.H. A Neural Implementation of the Kalman Filter. Journal of Advanced Research in Dynamical and Control Systems 12, 1513–1519 (2020).

29. Xivry, J.O.d., Coppe, S., Blohm, G. & Lefèvre, P. Kalman filtering naturally accounts for visually guided and predictive smooth pursuit dynamics. The Journal of Neuroscience 33, 17301–17313 (2013).

30. Izawa, J., Rane, T., Donchin, O. & Shadmehr, R. Motor adaptation as a process of reoptimization. The Journal of Neuroscience 28, 2883–2891 (2008).

31. III, A.R.P., Mathys, C. & Corlett, P.R. Pavlovian conditioning-induced hallucinations result from overweighting of perceptual priors. Cognitive Neuropsychiatry 22, 453–460 (2017).

32. Donaldson, K.R., Novak, K.D., Foti, D., Marder, M., Perlman, G., Kotov, R. & Mohanty, A. Mismatch negativity deficits are associated with auditory hallucinations transdiagnostically across psychotic disorders. Journal of Abnormal Psychology 129, 570–580 (2020).

33. Donaldson, K.R., Jonas, K., Foti, D., Larsen, E.M., Mohanty, A. & Kotov, R. Mismatch negativity predicts longitudinal worsening of auditory hallucinations in psychotic disorders. Psychological Medicine, (2022).

34. MS, T., EJ, A., T, B., E, A., A, S., B, W., P, C., SS, S. & SC, D. Reduced susceptibility to the Ebbinghaus and Müller-Lyer illusions in schizophrenia. Frontiers in Psychology 4, (2013).

35. DJ, K., J, H., PA, C., VA, C. & I, S. Visual illusion susceptibility in schizophrenia, bipolar disorder, and healthy controls. Psychonomic Bulletin & Review 24, 734–751 (2016).

36. OD, H., J, K., E, K., D, S., M, S., A, A. & S, K. Striatal presynaptic dopamine in schizophrenia, part II: meta-analysis of [18F/11C]-DOPA PET studies. Schizophrenia Bulletin 39, 33–42 (2012).

37. Jaeger., T. Ebbinghaus illusions: Size contrast or contour interaction phenomena?. Perception & Psychophysics 24, 337–342 (1978).

38. AE, P., C, B., C, K., RE, G. & RC, G. Actively paranoid patients with SZ over attribute anger to neutral faces. Schizophrenia Research 125, 174–178 (2011).

39. SH, S., NY, S., HL, W., W, C. & PA, G. Attribution bias in ultra-high risk for psychosis and first-episode SZ. Schizophrenia Research 118, 54–61 (2010).

40. Yu, A.J. & Dayan, P. Uncertainty, Neuromodulation, and Attention. Neuron 46, 681–692 (2005).

41. Howes, O.D. & Kapur, S. The nature of dopamine dysfunction in SZ and what this means for treatment. Archives of General Psychiatry 69, (2012).

42. Woo, T.W., Spencer, K. & McCarley, R.W. Gamma Oscillation Deficits and the Onset and Early Progression of SZ. Harvard Review of Psychiatry 18, 173–189 (2010).

43. Umbricht, D. & Krljes, S. Mismatch negativity in SZ: a meta-analysis. Schizophrenia Research 76, 1–23 (2005).

44. Erickson, M.A., Ruffle, A. & Gold, J.M. A meta-analysis of mismatch negativity in SZ: from clinical risk to disease specificity and progression. Biological Psychiatry 79, 980–987 (2016).

45. Kahneman, D. & Tversky, A. Prospect Theory: An Analysis of Decision under Risk. Econometrica 47, 263–291 (1979).

46. Lines, J., Martin, E.D., Kofuji, P., Aguilar, J. & Araque, A. Astrocytes modulate sensory-evoked network activity. Nature Communications 11, 3689 (2020).

47. Paukert, M., Agarwal, A., Cha, J., Doze, V.A., Kang, J.U. & Bergles, D.E. Norepinephrine controls astroglial responsiveness to local circuit activity. Neuron 82, 1263–1270 (2014).

48. Verkhratsky, A. & Parpura, V. Astrocytes modulate behavior through calcium signaling. Neuroglia, 320–332 (2012).

49. Corkrum, M. et al. Dopamine-Evoked Synaptic Regulation in the Nucleus Accumbens Requires Astrocyte Activity. Neuron 105, 1036–1047 (2020).

50. Khakh, B.S. & Sofroniew, M.V. Diversity and roles of astrocyte calcium signals in behavior and cognition. Nature Neuroscience 13, 759–766 (2010).

51. Denève, S., Duhamel, J. & Pouget, A. Optimal Sensorimotor Integration in Recurrent Cortical Networks. The Journal of Neuroscience 27, 5744–5756 (2007).

52. Beck, J.M., Latham, P.E. & Pouget, A. Marginalization in Neural Circuits with Divisive Normalization. The Journal of Neuroscience 31, 15310–15319 (2011).

53. Denève, S. & Jardri, R. Circular inference: mistaken belief, misplaced trust. Current Opinion in Behavioral Sciences 11, 40–48 (2016).

54. Adams, R.A., Stephan, K.E., Brown, H.R., Frith, C.D. & Friston, K.J. The computational anatomy of psychosis. Frontiers in Psychiatry 4, (2013).

55. Sterzer, P. et al. The Predictive Coding Account of Psychosis. Biological Psychiatry 84, 634–643 (2018).

56. Garety, P.A., Gittins, M., Jolley, S., Bebbington, P., Dunn, G., Fowler, D., Freeman, D. & Kuipers, E. Differences in Cognitive and Emotional Processes Between Persecutory and Grandiose Delusions. Schizophrenia Bulletin 39, 629–639 (2012).

57. Olshausen, B.A. & Field, D.J. Emergence of simple-cell receptive field properties by learning a sparse code for natural images. Nature 381, 607–609 (1996).

58. Ruderman, D.L. & Bialek, W. Statistics of natural images: Scaling in the woods. Physical Review Letters 73, 814–817 (1994).

59. Voss, R.F. & Clarke, J. 1/f noise” in music and speech. Nature 258, 317–318 (1975).

60. Baumeister, R.F., Bratslavsky, E., Finkenauer, C. & Vohs, K.D. Bad is stronger than good. Review of General Psychology 5, 323–370 (2001).

61. Dodds, P.S. et al. Human language reveals a universal positivity bias. Proceedings of the National Academy of Sciences 112, 2389–2394 (2015).

62. Boucher, J. & Osgood, C.E. The Pollyanna hypothesis. Journal of Verbal Learning and Verbal Behavior 8, 1–8 (1969).

63. Peter, V., McArthur, G. & Thompson, W.F. Effect of deviance direction and calculation method on duration and frequency mismatch negativity (MMN). Neuroscience Letters 482, 71–75 (2010).

64. Macdonald, M. & Campbell, K. Event-Related Potential Measures of a Violation of an Expected Increase and Decrease in Intensity. PLoS ONE 8, e76897 (2013).

65. Leung, S., Croft, P.G., Michie, P.T., Schall, U., Croft, R.J., O’Neill, D. & Todd, J. Acute glycine administration increases mismatch negativity in chronic schizophrenia. Frontiers in Human Neuroscience 9, (2015).

66. Kantrowitz, J.T., Malhotra, N., Hart, J., Huber, M., Apter, M.L. & Javitt, D.C. High dose D-serine in the treatment of schizophrenia. Schizophrenia Research 121, 125–130 (2010).

67. Lane, H., Lin, C., Huang, Y., Liao, C., Chang, Y. & Tsai, G.E. A randomized, double-blind, placebo-controlled comparison study of sarcosine (N-methylglycine) and D-serine add-on treatment for schizophrenia. The International Journal of Neuropsychopharmacology 13, 451 (2009).

68. Carter, C.S. et al. Functional hypofrontality and working memory dysfunction in schizophrenia. American Journal of Psychiatry 155, 1285–1287 (1998).

69. Milad, M.R. & Quirk, G.J. Fear Extinction as a Model for Translational Neuroscience. Annual Review of Psychology 63, 129–151 (2012).

70. Monosov, I.E. Anterior cingulate is a source of valence-specific information about value and uncertainty. Nature Communications 8, 134 (2017).

71. McHugh, S.B. et al. Aversive prediction error signals in the amygdala. Journal of Neuroscience 34, 9024–9033 (2014).

