## Supplemental Material for "A Dual-Pathway Prediction Error Model of Schizophrenia Spectrum Disorders: Bridging NMDA Hypofunction and Dopaminergic Hyperfunction"

```
"""
Supplemental Material: Code for Figure 2
==========================================

Description:
    This script generates the panels for Figure 2, illustrating the differences in
    posterior inference and prediction errors between Control and SSD conditions
    during 'Omission' and 'Presence' tasks.

    - Figure 2c: Posterior Estimates (M) under the Omission condition.
    - Figure 2d: Prediction Error magnitude (MMN) for both Omission and Presence.

    The simulation implements a simplified 1D Kalman Filter step.

Dependencies:
    - numpy
    - matplotlib

"""

import numpy as np
import matplotlib.pyplot as plt
import os
from matplotlib.lines import Line2D

#
==========================================================================
=======
# Configuration & Parameters
#
==========================================================================
=======

# Use a headless backend if no interactive backend is available (for server runs)
if os.environ.get("MPLBACKEND") is None:
    plt.switch_backend('Agg')

# Model Parameters
K_VALUE = 0.618       # Steady-state Kalman Gain
ALPHA_CONTROL = 1.0   # Attenuation factor for negative PE (Control)
ALPHA_DISEASE = 0.6   # Attenuation factor for negative PE (SSD - NMDA hypofunction)
A_PARAM = 0.9         # Autoregressive parameter

# Plotting Style
plt.rcParams['font.family'] = 'serif'
plt.rcParams['font.serif'] = ['Times New Roman']
plt.rcParams['font.size'] = 14
plt.rcParams['axes.linewidth'] = 1.2
```

```
plt.rcParams['xtick.major.width'] = 1.2
plt.rcParams['ytick.major.width'] = 1.2

#
==========================================================================
=======
# Simulation Logic
#
==========================================================================
=======

def run_simulation(alpha, prior_val, obs_val):
    """
    Simulates a single time-step update of the latent state.

    Args:
        alpha (float): Scaling factor for negative prediction error (e.g., 1.0 or 0.6).
        prior_val (float): The prior belief state (M_t-1).
        obs_val (float): The current observation (o_t).

    Returns:
        tuple: (Posterior Estimate, Effective Total Prediction Error)
    """
    # Prediction: M_pred = a * M_t-1
    prediction = A_PARAM * prior_val
    obs = obs_val

    # Calculate Raw Prediction Errors (Rectified)
    if obs >= prediction:
        e_plus = obs - prediction
        e_minus = 0.0
    else:
        e_plus = 0.0
        e_minus = prediction - obs

    # Apply bias (alpha) to the negative error component
    effective_e_minus = alpha * e_minus
    effective_pe_total = e_plus + effective_e_minus

    # Update: M_post = M_pred + K * (E+ - alpha * E-)
    update = K_VALUE * (e_plus - effective_e_minus)
    posterior = prediction + update

    return posterior, effective_pe_total

#
==========================================================================
=======
```

```
# Main Execution
#
==========================================================================
=======

if __name__ == "__main__":

    # --- Condition 1: Omission (Prior=1, Obs=0) ---
    # Simulates Hallucination-like state where input is absent but expected.
    prior_hallucination = 1.0
    obs_hallucination = 0.0

    post_omit_ctrl, mmn_omit_ctrl = run_simulation(ALPHA_CONTROL, prior_hallucination,
obs_hallucination)
    post_omit_ssd,  mmn_omit_ssd  = run_simulation(ALPHA_DISEASE, prior_hallucination,
obs_hallucination)

    # --- Condition 2: Presence (Prior=0, Obs=1) ---
    # Simulates standard perception where input is present but unexpected.
    prior_presence = 0.0
    obs_presence = 1.0

    post_pres_ctrl, mmn_pres_ctrl = run_simulation(ALPHA_CONTROL, prior_presence,
obs_presence)
    post_pres_ssd,  mmn_pres_ssd  = run_simulation(ALPHA_DISEASE, prior_presence,
obs_presence)


    #
==========================================================================
=======
    # Plotting Figure 2c: Posterior Estimates
    #
==========================================================================
=======
    labels_legend = ['Control', 'SSD']

    fig1, ax1 = plt.subplots(figsize=(6, 6))

    x_c = 0
    width = 0.35

    ctrl_posts = [post_omit_ctrl]
    ssd_posts = [post_omit_ssd]

    rects1 = ax1.bar(x_c - width/2, ctrl_posts, width, label='Control', color='#808080',
edgecolor='black', alpha=0.9)
```

```
    rects2 = ax1.bar(x_c + width/2, ssd_posts, width, label='SSD', color='#800080',
edgecolor='black', alpha=0.9)

    ax1.set_ylabel('Posterior Estimate (M)', fontsize=14)
    ax1.set_xticks([x_c - width/2, x_c + width/2])
    ax1.set_xticklabels(['Control', 'SSD'], fontsize=12)
    ax1.grid(axis='y', linestyle=':', alpha=0.4)
    ax1.set_ylim(-0.1, 1.3)

    # Add reference lines for Prior and Observation
    w_line = 0.35 * 2 + 0.1
    ax1.plot([x_c - w_line/2, x_c + w_line/2], [1.0, 1.0], color='blue', linestyle='--',
linewidth=1.5) # Prior
    ax1.plot([x_c - w_line/2, x_c + w_line/2], [0.0, 0.0], color='gray', linestyle='--',
linewidth=1.5) # Observation

    # Custom Legend
    legend_elements = [
        Line2D([0], [0], color='blue', linestyle='--', lw=1.5, label='Prior'),
        Line2D([0], [0], color='gray', linestyle='--', lw=1.5, label='Observation')
    ]
    ax1.legend(handles=legend_elements, loc='upper right', frameon=False, fontsize=10)

    ax1.bar_label(rects1, fmt='%.2f', padding=3)
    ax1.bar_label(rects2, fmt='%.2f', padding=3)

    plt.tight_layout()
    fig1.savefig('Figure_2c.png', dpi=300, bbox_inches='tight')
    print("Saved Figure_2c.png")


    #
===========================================================================
    # Plotting Figure 2d: Prediction Error / MMN
    #
===========================================================================
    fig2, ax2 = plt.subplots(figsize=(6, 6))

    x_d = np.arange(2) # Groups: Omission, Presence

    ctrl_mmns = [mmn_omit_ctrl, mmn_pres_ctrl]
    ssd_mmns = [mmn_omit_ssd, mmn_pres_ssd]

    rects3 = ax2.bar(x_d - width/2, ctrl_mmns, width, label='Control', color='#808080',
edgecolor='black', alpha=0.9)
```

```
    rects4 = ax2.bar(x_d + width/2, ssd_mmns, width, label='SSD', color='#800080',
edgecolor='black', alpha=0.9)

    ax2.set_ylabel('Prediction Error Strength', fontsize=14)
    ax2.set_xticks(x_d)
    ax2.set_xticklabels(['Omission', 'Presence'], fontsize=12)
    ax2.legend()
    ax2.grid(axis='y', linestyle=':', alpha=0.4)
    ax2.set_ylim(0, 1.3)

    ax2.bar_label(rects3, fmt='%.2f', padding=3)
    ax2.bar_label(rects4, fmt='%.2f', padding=3)

    plt.tight_layout()
    fig2.savefig('Figure_2d.png', dpi=300, bbox_inches='tight')
    print("Saved Figure_2d.png")
```

```
"""
Supplemental Material: Code for Figure 3
==========================================

Description:
    This script generates Figure 3c, which simulates the Ebbinghaus illusion.
    It demonstrates how dopaminergic hyperfunction (increased gain beta) leads to
    a reduction in the illusion magnitude by overweighting sensory evidence
    relative to contextual priors.

    - Condition 1: Large Flankers (Shrinking Context), Prior < Observation.
    - Condition 2: Small Flankers (Enlarging Context), Prior > Observation.

Dependencies:
    - numpy
    - matplotlib

"""

import numpy as np
import matplotlib.pyplot as plt
import os

#
==========================================================================
=======
# Configuration & Parameters
#
==========================================================================
=======

if os.environ.get("MPLBACKEND") is None:
    plt.switch_backend('Agg')

# Plotting Style
plt.rcParams['font.family'] = 'serif'
plt.rcParams['font.serif'] = ['Times New Roman']
plt.rcParams['font.size'] = 14
plt.rcParams['axes.linewidth'] = 1.2
plt.rcParams['xtick.major.width'] = 1.2
plt.rcParams['ytick.major.width'] = 1.2

# Model Parameters
A_PARAM = 0.9
C_PARAM = 1.0
B_PARAM = np.sqrt(0.1) # Process noise standard deviation (Q = b^2)
D_PARAM = 1.0          # Observation noise standard deviation (R = d^2)
```

```
Q_VAL = B_PARAM**2
R_VAL = D_PARAM**2

OBS_VAL = 2.0  # Veridical Object Size

#
==========================================================================
=======
# Helper Functions
#
==========================================================================
=======

def get_steady_state_K(a, c, q, r):
    """
    Computes the steady-state Kalman Gain (K) via iteration.

    Args:
        a, c (float): System matrices/coefficients.
        q, r (float): Process and observation noise variances.

    Returns:
        float: Converged Kalman Gain.
    """
    p = 1.0
    for _ in range(1000):
        # Time update
        p_pred = a**2 * p + q
        # Kalman Gain
        k = p_pred * c / (c**2 * p_pred + r)
        # Measurement update
        p_new = (1 - k * c) * p_pred

        if abs(p - p_new) < 1e-9:
            break
        p = p_new
    return k

# Pre-calculate Optimal Gain
K_OPTIMAL = get_steady_state_K(A_PARAM, C_PARAM, Q_VAL, R_VAL)


def run_simulation(beta, prior_mean):
    """
    Simulates the specific Ebbinghaus illusion trial.

    Args:
        beta (float): Dopamine gain factor multiplying the Kalman Gain.
```

```
                beta=1.0 -> Optimal/Control.
                beta>1.0 -> Overweighted sensory evidence (SSD).
        prior_mean (float): The contextual prior (M_t-1).

    Returns:
        float: The posterior size estimate.
    """
    # Prediction Step based on context
    prediction = A_PARAM * prior_mean

    # Modulated Kalman Gain
    k_mod = beta * K_OPTIMAL

    # Update Step: Posterior = Prediction + K * PredictionError
    posterior = prediction + k_mod * (OBS_VAL - prediction)
    return posterior

#
==========================================================================
=======
# Main Execution
#
==========================================================================
=======

if __name__ == "__main__":

    fig, ax = plt.subplots(figsize=(8, 6))

    # Simulation Settings
    beta_control = 1.0
    beta_ssd = 1.15

    # --- Condition 1: Large Flankers (Shrinking Context) ---
    # The context implies the object is small (Prior=1.0), while true size is 2.0.
    prior_shrink = 1.0
    val_shrink_ctrl = run_simulation(beta_control, prior_shrink)
    val_shrink_ssd  = run_simulation(beta_ssd, prior_shrink)

    # --- Condition 2: Small Flankers (Enlarging Context) ---
    # The context implies the object is large (Prior=3.0), while true size is 2.0.
    prior_enlarge = 3.0
    val_enlarge_ctrl = run_simulation(beta_control, prior_enlarge)
    val_enlarge_ssd  = run_simulation(beta_ssd, prior_enlarge)

    # Prepare Data for Plotting
    labels = ['Large Flankers\n(Shrinking Context)', 'Small Flankers\n(Enlarging Context)']
    control_means = [val_shrink_ctrl, val_enlarge_ctrl]
```

```
    ssd_means = [val_shrink_ssd, val_enlarge_ssd]

    x = np.arange(len(labels))
    width = 0.35

    # Draw Bars
    rects1 = ax.bar(x - width/2, control_means, width, label='Control ($\\beta=1.0$)',
color='gray', alpha=0.8, edgecolor='k')
    rects2 = ax.bar(x + width/2, ssd_means, width, label='SSD ($\\beta=1.15$)', color='purple',
alpha=0.8, edgecolor='k')

    # --- Reference Lines ---
    # 1. True Observation
    ax.axhline(OBS_VAL, color='green', linestyle='-', linewidth=2, label='Observation (True
Size)', zorder=0)

    # 2. Contextual Priors
    # Large Flankers (Index 0)
    ax.hlines(prior_shrink, x[0]-0.45, x[0]+0.45, colors='blue', linestyles='--', linewidth=2,
label='Context Prior (Biased)')
    ax.text(x[0], prior_shrink - 0.25, 'Prior = 1.0', ha='center', color='blue', fontsize=12)

    # Small Flankers (Index 1)
    ax.hlines(prior_enlarge, x[1]-0.45, x[1]+0.45, colors='blue', linestyles='--', linewidth=2)
    ax.text(x[1], prior_enlarge + 0.05, 'Prior = 3.0', ha='center', color='blue', fontsize=12)

    # Formatting
    ax.set_ylabel('Estimated Size (Posterior)', fontsize=14)
    ax.set_xticks(x)
    ax.set_xticklabels(labels, fontsize=13)
    ax.set_ylim(0.5, 3.0)

    ax.legend(loc='upper center', bbox_to_anchor=(0.5, -0.15), ncol=2, fontsize=12)
    ax.bar_label(rects1, fmt='%.3f', padding=3, fontsize=10)
    ax.bar_label(rects2, fmt='%.3f', padding=3, fontsize=10)
    ax.grid(axis='y', linestyle='--', alpha=0.3)

    plt.tight_layout()
    output_filename = "Figure_3c_SSD.png"
    plt.savefig(output_filename, dpi=300, bbox_inches='tight')
    print(f"Saved {output_filename}")
```

```
"""
Supplemental Material: Code for Figure 4
==========================================

Description:
    This script simulates the temporal dynamics of the Dual-Pathway Model for SSD.
    It generates the time-series plots (Figure 4) showing how interactions between
    Dopaminergic hyperfunction and NMDA receptor hypofunction lead to distinct
    patterns of belief updating and instability.

    - Panel A: Example trajectories of True State vs. Observation.
    - Panel B: Estimation errors for Control, SSD (Combined), Dopamine-only,
               and NMDA-only conditions.
    - Panel C: Root Mean Square of Successive Differences (RMSSD) quantifying
               belief instability.
    - Panel D: Heatmap of RMSSD across the parameter space (Dopamine x NMDA).

Dependencies:
    - numpy
    - matplotlib

"""

import numpy as np
import os
import sys
import matplotlib as mpl
import matplotlib.pyplot as plt
from pathlib import Path


#
==========================================================================
=======
# Configuration & Utilities
#
==========================================================================
=======

def _prefer_interactive_backend():
    """Sets an interactive backend if available, otherwise falls back to Agg."""
    if os.environ.get('MPLBACKEND'):
        return
    try:
        current = mpl.get_backend().lower()
    except Exception:
        current = ''
    if 'agg' in current or current == '':
```

```
    candidates = ['MacOSX', 'TkAgg'] if sys.platform == 'darwin' else ['TkAgg', 'Qt5Agg',
'QtAgg']
    for cand in candidates:
        try:
            mpl.use(cand, force=True)
            break
        except Exception:
            continue

_prefer_interactive_backend()


def configure_matplotlib_for_paper():
    """Configures matplotlib parameters for publication-quality figures."""
    mpl.rcParams.update({
        'figure.dpi': 110,
        'savefig.dpi': 300,
        'font.family': 'serif',
        'font.serif': ['Times New Roman'],
        'font.size': 14,
        'axes.titlesize': 15,
        'axes.labelsize': 14,
        'axes.linewidth': 1.2,
        'axes.titlepad': 10.0,
        'xtick.labelsize': 13,
        'ytick.labelsize': 13,
        'xtick.major.width': 1.2,
        'ytick.major.width': 1.2,
        'xtick.direction': 'out',
        'ytick.direction': 'out',
        'grid.alpha': 0.3,
        'lines.linewidth': 2.0,
        'legend.frameon': True,
        'legend.fontsize': 12,
    })

def beautify_ax(ax):
    """Removing top and right spines for a cleaner look."""
    ax.spines['top'].set_visible(False)
    ax.spines['right'].set_visible(False)
    ax.grid(True, alpha=0.25)
    ax.tick_params(length=4, width=1)


#
===========================================================================
=======
# Kalman Filter Class
```

```
# 
=======================================================================
=======

class KalmanFilter:
    def __init__(self, a, c, b, d, s0, P0, alpha, beta):
        """
        Initializes the Kalman Filter with dual-pathway modulation.

        Args:
            a, c (float): System transition and observation matrices (scalars).
            b, d (float): Noise standard deviations for process and observation.
            s0, P0 (float): Initial state and covariance.
            alpha (float): Inhibitory pathway gain (NMDA modulation on negative error channel).
            beta (float): Dopamine gain factor applied to the Kalman Gain.
        """
        self.A = a
        self.B = c
        self.Q = b**2  # Process noise variance
        self.R = d**2  # Observation noise variance

        self.s = s0
        self.P = P0
        self._prev_s = self.s

        # Dual-pathway parameters
        self.alpha = alpha  # Modulates sensitivity to negative prediction errors (NMDA)
        self.beta = beta    # Dopaminergic Gain

        self.err = []
        self.state = []

    def step(self, o):
        # 1. Prediction Step
        s_pred = self.A * self.s
        P_pred = self.A * self.P * self.A + self.Q

        # 2. Kalman Gain Computation
        K = P_pred * self.B / (self.B * P_pred * self.B + self.R)

        # 3. Covariance Update
        self.P = (1 - K * self.B) * P_pred

        # 4. Observation Prediction
        obs_pred = self.B * s_pred

        # 5. Dual-Pathway Prediction Error Calculation
        # E+ (Positive Error): Input exceeds prediction
```

```
        # E- (Negative Error): Prediction exceeds input
        E_pos = max(0, o - obs_pred)
        E_neg = max(0, -o + obs_pred)

        # 6. State Update with Modulated Correction
        # Paper Equation: M_t = a*M_{t-1} + beta * K * (E_pos - alpha * E_neg)
        correction = self.beta * K * (E_pos - self.alpha * E_neg)
        self.s = s_pred + correction

        self.err.append(self.s - self._prev_s)
        self.state.append(self.s)
        self._prev_s = self.s


#
=========================================================================
=======
# Main Simulation Execution
#
=========================================================================
=======

if __name__ == "__main__":
    configure_matplotlib_for_paper()

    # --- Simulation Parameters ---
    T, n_runs = 30, 1

    # Fixed seed for reproducibility (Okabe-Ito color palette friendly)
    rng_master = np.random.default_rng(102)

    a, c = 0.9, 1.0
    b, d = np.sqrt(0.1), 1.0  # Noise StDevs
    s0, P0 = 0.0, 1.0

    # Okabe-Ito Color Palette for accessibility
    okabe_ito = {
        'black':   '#000000',
        'orange':  '#E69F00',
        'skyblue': '#56B4E9',
        'green':   '#009E73',
        'yellow':  '#F0E442',
        'blue':    '#0072B2',
        'verm':    '#D55E00',
        'purple':  '#CC79A7',
    }
```

```
    # Model Conditions (e=1.0 is normal NMDA, i=0.6 is NMDA hypofunction on interneurons
leading to disinhibition/attenuation issues logic per paper)
    # Note: 'i' in code maps to 'e' in paper text? Let's assume the parameters e/i correspond to
the paper's alpha_plus/alpha_minus logic.
    # Code mapping:
    #   N-condition (Control): e=1, i=1, beta=1
    #   S-condition (Synergy/SSD): e=1, i=0.6, beta=1.15
    models = {
        'N-condition': dict(e=1.0, i=1.0, beta=1.0,  color=okabe_ito['black']),   # Control
        'S-condition': dict(e=1.0, i=0.6, beta=1.15,  color='purple'),           # SSD (Both)
        'A-condition': dict(e=1.0, i=1.0, beta=1.15,  color='red'),             # Dopamine
Hyperfunction Only
        'P-condition': dict(e=1.0, i=0.6, beta=1.0,  color='blue'),             # NMDA Hypofunction
Only
    }

    # Data Containers
    true_states  = np.zeros((n_runs, T))
    observations = np.zeros((n_runs, T))
    estimates = {k: np.zeros((n_runs, T)) for k in models}
    errors    = {k: np.zeros((n_runs, T)) for k in models}
    rmssd     = {k: np.zeros(n_runs)      for k in models}

    # --- Run Simulation ---
    for r in range(n_runs):
        rng = np.random.default_rng(rng_master.integers(2**32))

        # Generate Episode
        s_true = [0.0]
        obs = []
        for _ in range(T):
            s_next = a * s_true[-1] + rng.normal(0, b)
            obs.append(s_next + rng.normal(0, d))
            s_true.append(s_next)
        true_states[r]  = s_true[1:]
        observations[r] = obs

            # Run Filters
        kf = {k: KalmanFilter(a, c, b, d, s0, P0, cfg['alpha'], cfg['beta'])
              for k, cfg in models.items()}

        for k in kf:
            for o in obs:
                kf[k].step(o)

        # Store Results
        for k in models:
            estimates[k][r] = kf[k].state
```

```
        errors[k][r]    = np.array(kf[k].state) - np.array(s_true[1:])
        rmssd[k][r]     = np.sqrt(np.mean(np.diff(kf[k].err)**2))

    ts = np.arange(1, T+1)


    #
==========================================================================
=======
    # Panel A: True State & Observation
    #
==========================================================================
=======
    figA, axA = plt.subplots(figsize=(8.2, 4.2))
    true_state_run = true_states[0]
    obs_run        = observations[0]

    axA.plot(ts, true_state_run, '-', color=okabe_ito['black'], lw=2.2, label='True state')
    axA.plot(ts, obs_run,  '-', color='gray', lw=2.0, label='Observation')

    axA.set_xlim(1, T)
    axA.set_xlabel('Time step')
    axA.set_ylabel('Social Value (Hostility)')
    axA.legend(loc='upper left', bbox_to_anchor=(1.05, 1.0))
    beautify_ax(axA)


    #
==========================================================================
=======
    # Panel B: Estimation Error Dynamics
    #
==========================================================================
=======
    figB, axes = plt.subplots(3,1,figsize=(8.8, 9.2), sharex=True)
    pairs=[
        ('S-condition', 'Control vs Both'),
        ('A-condition', 'Control vs Dopamine hyperfunction'),
        ('P-condition', 'Control vs NMDA receptor hypofunction')
    ]

    label_map = {
        'N-condition': 'Control',
        'S-condition': 'Both',
        'A-condition': 'Dopamine hyperfunction',
        'P-condition': 'NMDA receptor hypofunction'
    }
```

```
for ax, (d_key, title) in zip(axes, pairs):
    ymax_abs = 0
    for key in ['N-condition', d_key]:
        err_run = errors[key][0]
        ax.plot(ts, err_run, color=models[key]['color'], label=label_map.get(key, key))
        ymax_abs = max(ymax_abs, np.max(np.abs(err_run)))

    ax.axhline(0, color='#999999', lw=1.2, ls='--', zorder=0)

    margin = 0.05
    lim = np.ceil((ymax_abs + margin) * 10) / 10
    ax.set_ylim(-lim, lim)
    ax.set_ylabel('Estimation error')
    beautify_ax(ax)
    ax.legend(loc='upper left', bbox_to_anchor=(1.05, 1.0))

axes[-1].set_xlabel('Time step')


#
==========================================================================
=======
# Panel C: RMSSD Boxplot
#
==========================================================================
=======
figC, axC = plt.subplots(figsize=(6.6, 4.4))
order = list(models.keys())
data  = [rmssd[k] for k in order]
colors_list = [models[k]['color'] for k in order]

bp = axC.boxplot(
    data,
    tick_labels=[label_map.get(k, k).replace(' ', '\n') for k in order],
    patch_artist=True,
    showmeans=False,
    boxprops=dict(linewidth=1.2, color='black'),
    whiskerprops=dict(linewidth=1.2, color='black'),
    capprops=dict(linewidth=1.2, color='black'),
    medianprops=dict(color='none'),
)

for patch in bp['boxes']:
    patch.set_facecolor('white')
    patch.set_edgecolor('black')

for i, line in enumerate(bp['medians']):
    x_data = line.get_xdata()
```

```
        y_val_med = line.get_ydata()[0]
        color_val = colors_list[i]

        # Dashed median line
        axC.plot(x_data, [y_val_med, y_val_med], color=color_val, linestyle='--', linewidth=2.0,
zorder=3)
        # Triangles at ends
        axC.plot(x_data[0], y_val_med, marker='^', color='black', markersize=8, zorder=4)
        axC.plot(x_data[1], y_val_med, marker='^', color='black', markersize=8, zorder=4)

    axC.yaxis.grid(True, linestyle='-', alpha=0.5)
    axC.set_ylabel('Root Mean Square (Instability)')


    #
=========================================================================
=======
    # Panel D: Heatmap (Dopamine x NMDA)
    #
=========================================================================
=======
    Beta_vals = np.linspace(1.0, 1.5, 21)
    Alpha_vals = np.linspace(1.0, 0.4, 21)   # Sweep NMDA integrity (alpha) down to 0.4
    heat = np.zeros((len(Beta_vals), len(Alpha_vals)))

    for r_idx in range(len(Beta_vals)):
        for c_idx in range(len(Alpha_vals)):
            b_val = Beta_vals[r_idx]
            a_val = Alpha_vals[c_idx]
            rms_list = []

            # Using the same fixed observations for consistency
            for obs in observations:
                flt = KalmanFilter(a, c, b, d, s0, P0, alpha=a_val, beta=b_val)
                for o in obs:
                    flt.step(o)
                rms_list.append(np.sqrt(np.mean(np.diff(flt.err)**2)))
            heat[r_idx, c_idx] = rms_list[0]

    figH, axH = plt.subplots(figsize=(7.0, 5.4))
    im = axH.imshow(
        heat,
        origin='lower',
        aspect='auto',
        interpolation='nearest',
        extent=[Alpha_vals.max(), Alpha_vals.min(), Beta_vals.min(), Beta_vals.max()],
        cmap=mpl.colors.LinearSegmentedColormap.from_list('rpb', ['blue', 'purple', 'red'])
    )
```

```
    axH.set_xlabel('NMDA bias (alpha)')
    axH.set_ylabel('Dopamine gain')
    cbar = figH.colorbar(im, ax=axH, label='RMSSD')
    beautify_ax(axH)


    #
==========================================================================
=======
    # Save Figures
    #
==========================================================================
=======
    try:
        base_dir = Path(os.getcwd())
        # Renamed output files to match Figure 4 convention
        for fig, stem in [(figA,'Figure_4A'), (figB,'Figure_4B'), (figC,'Figure_4C'),
(figH,'Figure_4D')]:
            png_path = base_dir / f'{stem}.png'
            fig.savefig(png_path.as_posix(), dpi=300, bbox_inches='tight')
            print(f"Saved {png_path.name}")
    except Exception as e:
        print(f"Error saving figures: {e}")
```
